# Intratumoral cDC1-T Cell Clusters Serve as Sites of Local Costimulation to Enhance CTL-Mediated Tumor Rejection

**DOI:** 10.1101/2025.05.19.654765

**Authors:** Catherine B. Carbone, Markus Brown, Camille-Charlotte Balança, Anthony C. Antonelli, Shiuh-Ming Luoh, Karl L. Banta, Vincent Rouilly, Ximo Pechuan-Jorge, Guadalupe Ortiz-Muñoz, Jeanne Cheung, Ines Marin, Ellen Duong, Aditya A. Anand, Caleb K. Chan, Anugraha Rajagopalan, Justin T. Gibson, Amanda Lorentzian, Elisa Penna, Jian Jiang, Linda Rangell, Yuxin Liang, Jérémie Decalf, Rosa Barreira da Silva, Ying Zhu, Shannon J. Turley, Geraldine A. Strasser, Sandra Rost, Shomyseh Sanjabi, Kathryn Geiger-Schuller, Lélia Delamarre, Ira Mellman, Christine Moussion

## Abstract

T cells are essential for anti-tumor immunity, but their ability to eliminate tumors depends on coordinated interactions with type 1 conventional dendritic cells (cDC1s). While cDC1s are known for cross-presenting tumor-derived antigens in lymph nodes to prime CD8+ T cells, their role within the tumor itself remains less well understood. Here, we use the Skin Tumor Array by Micro-Poration (STAMP) model to investigate how cDC1-T cell interactions shape immune responses and influence tumor fate. Our data reveal that it is the spatial distribution of both cDC1s and T cells that determines whether a tumor can be rejected. We defined three primary immunotypes based on the spatial distribution of T cells and cDC1s: T cell-inflamed/dendritic cell-inflamed (TC-In/DC-In) tumors, where T cells and cDC1s co-infiltrate the tumor; T cell-inflamed/dendritic cell-excluded (TC-In/DC-Ex) tumors, where T cells infiltrate but cDC1s remain at the periphery; and T cell-excluded/dendritic cell-excluded (TC-Ex/DC-Ex) tumors, which lack both cDC1 and T cell infiltration. Notably, TC-In/DC-In tumors are more likely to undergo rejection, whereas TC-In/DC-Ex tumors persist despite T cell infiltration. Within TC-In/DC-In tumors, cDC1s engage in direct interactions with T cells, upregulate co-stimulatory molecules, and sustain effector T cell responses, while cDC1s in TC-In/DC-Ex tumors express higher migration-associated genes, suggesting a propensity to exit the tumor. We further show that chemokine modulation, particularly through CXCL9, CCL5, and XCL1, can reshape immune infiltration patterns to promote intra-tumoral cDC1-T cell clustering and improve tumor rejection. These findings underscore the unexpectedly important role of cDC1 positioning and function in sustaining effective anti-tumor immunity and highlight spatially organized cDC1-T cell clusters as critical hubs for local T cell activation.

## Introduction

The spatial distribution of T cells within a tumor (immunotype) is a strong predictor of immunotherapy response—tumors where T cells are restricted to the stroma (excluded) or absent (desert) are often resistant to treatment, whereas tumors with abundant intratumoral T cells (inflamed) are typically associated with better outcomes (Chen et al., 2017). However, not all inflamed tumors respond to therapy, indicating that T cell presence alone is insufficient to predict patient responses (Mellman et al., 2013). Additional factors, including the composition of the tumor microenvironment (TME), may determine whether immune responses are productive or dysfunctional. Among these factors, antigen-presenting cells (APCs) play a critical role in shaping anti-tumor immunity, with increasing evidence pointing to type 1 conventional dendritic cells (cDC1s) as key regulators of T cell-mediated tumor rejection. cDC1s are essential for priming CD8+ T cells in lymph nodes (Hildner et al., 2008; Roberts et al., 2016), and studies suggest they also have direct functions within tumors (Broz et al 2014, Spranger et al., 2017), where they may support T cell activation, persistence, and effector function (Bottcher et al., 2018; Di Pilato et al., 2021). Within tumors, the functional state and spatial organization of cDC1s have been identified as key correlates of whether an inflamed tumor is rejected or persists (Magen & Hamon, 2023; Chen et al., 2024). However, these associations alone do not fully explain why some tumors fail to respond to treatment, highlighting the need for functional, time-resolved data and spatially resolved in vivo models to uncover how cDC1s sustain productive immune responses.

Dendritic cells (DCs) comprise a heterogeneous population that includes cDC1s, cDC2s, monocyte-derived DCs (moDCs), and plasmacytoid DCs (pDCs), each with distinct yet overlapping functions. Among them, cDC1s are uniquely specialized for antigen cross-presentation and are essential for priming CD8+ T cells and initiating antitumor immunity (Hildner et al., 2008; Roberts et al., 2016). While other APCs can assume antigen-presenting roles in specific contexts—such as inflammatory monocytes (Elewaut et al., 2024) or cDC2s (Duong et al., 2022, Prokhnevska et al, 2023) supporting CD8+ T cell activation in tumors — cDC1s remain the primary APCs responsible for initiating tumor-specific CD8+ T cell responses. Beyond the diversity of APCs, cDC1s themselves exhibit substantial variability in activation, maturation, and migratory states, all of which influence their ability to sustain immune responses and must be further understood.

Inside the tumor microenvironment, the distinct phenotypic states of cDC1s influence their ability to orchestrate immune activation and sustain anti-tumor responses. cDC1s have been ascribed roles in recruiting effector T cells through CXCL9 and CXCL10 secretion (Meiser et al., 2023; Bayerl et al., 2023), presenting antigens via MHC-I and MHC-II, and providing co-stimulatory signals such as CD80, CD86, CD40, OX40L, and 4-1BBL to sustain T cell activation (Lee et al., 2024; Ziblat et al., 2024; Ferris et al., 2021). They can also support T cell persistence through IL-12 production, which enhances CD8+ T cell differentiation (Magen & Hamon, 2023), and IL-15 trans-presentation, which promotes long-term survival of CD8+ effectors (Di Pilato et al., 2021).

Given the functional heterogeneity of cDC1s, a critical question is how their positioning in and around a tumor affects their ability to coordinate immunity. To explore this, we used STAMP (Skin Tumor Array by Micro-Poration), a high-throughput preclinical imaging technology in which tumor arrays are implanted in mouse ear pinnae, allowing individual tumors to be tracked longitudinally until they either regress or reach experimental endpoints (Ortiz-Muñoz et al., 2023). Conventional in vivo models and human studies require separate experiments to assess immune dynamics and tumor outcomes because capturing immune activity typically requires endpoint analyses or invasive sampling, which can disrupt the microenvironment. In contrast, STAMP enables non-invasive imaging of cDC1s, T cells, and tumor growth, directly linking immune cell positioning to tumor rejection within a single experiment.

Here, we leverage the STAMP model to uncover how the spatial organization and functional state of cDC1s within tumors shape T cell behavior and determine tumor fate. By combining live imaging, transcriptional profiling, and chemokine modulation, we identify spatially organized cDC1–T cell clusters as critical hubs of effector function and tumor rejection, providing a mechanistic explanation for why some T cell-inflamed tumors remain refractory to treatment.

## Results

### Imaging and Classification of cDC1/T Cell Interactions Reveals Distinct Immunotypes in Mouse and Human Tumors

The STAMP technique involves implanting genetically identical micro-tumors into an array of pores created in the skin of a mouse ear using an infrared laser device. (**Fig. 1A**). Despite being clonal, individual STAMP tumors exhibit distinct patterns of immune cell infiltration and distinct outcomes of rejection or progression (Ortiz-Muñoz et al Nature 2023). Since T cell priming occurs in a common draining lymph node shared by all tumors in the array, we can directly test how microenvironment-intrinsic interactions regulate tumor immunity between neighboring lesions.

**Figure 1.**
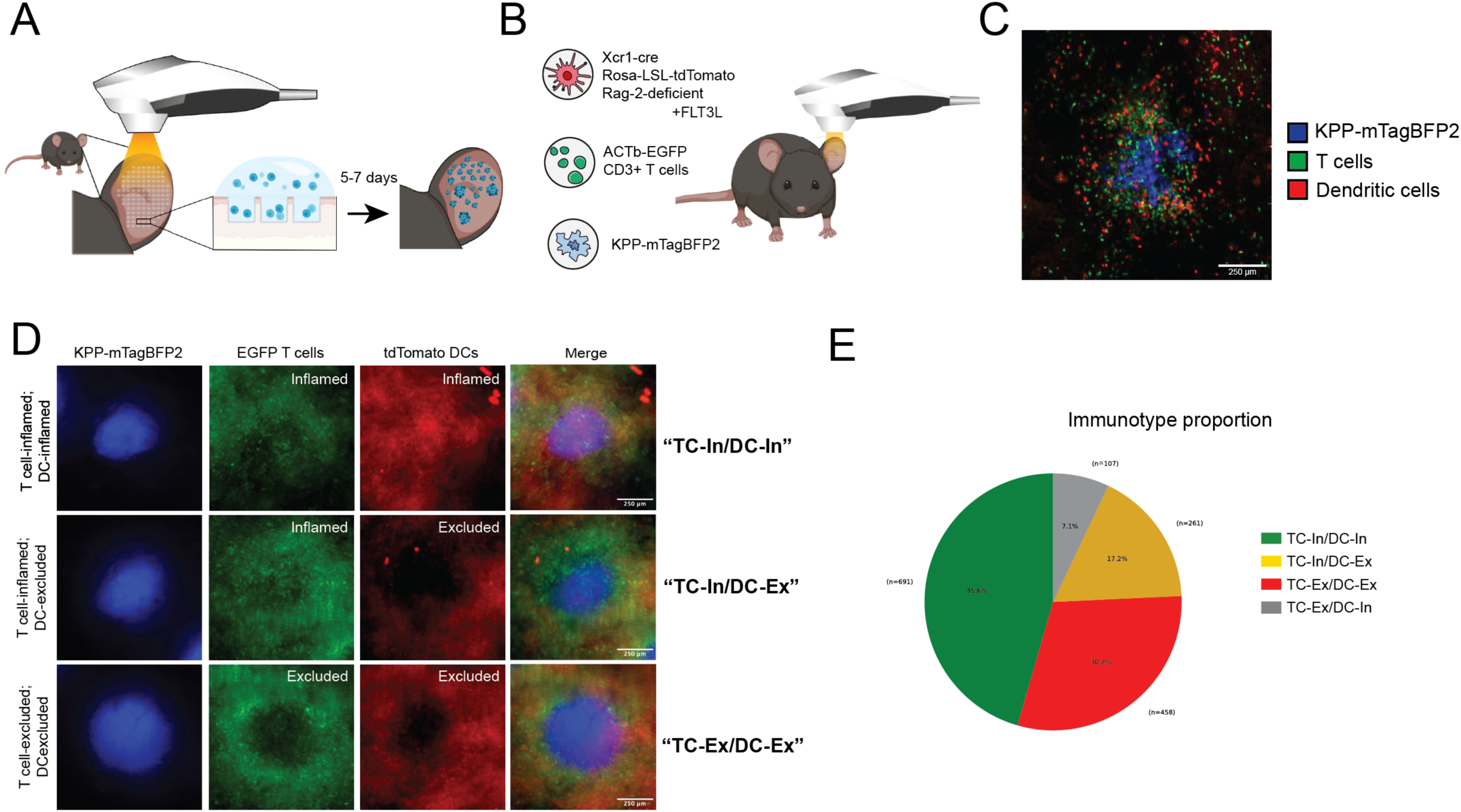
cDC1 and T cell localization defines distinct immunotypes in mouse and human tumors. (A) Schematic of STAMP tumor implantation procedure including formation of the micropore array, incubation with cell suspension, and resultant tumor array. (B) Genetic model for multiplexed imaging of cDC1s, T cells, and tumor cells in STAMP microtumor arrays. Xcr1-Cre; Rosa26-LSL-tdTomato mice specifically label cDC1s with tdTomato fluorescence. Crossing these mice to Rag2 KO mice eliminates endogenous T cells, while Flt3L treatment expands the cDC1 population for enhanced imaging. For STAMP experiments, EGFP-labeled T cells are adoptively transferred, and mTagBFP2+ tumor cells are implanted to establish tumors. (C) Representative 2-photon image of STAMP arrays showing KPP-mTagBFP2+ tumor cells (blue), tdTomato-labeled cDC1s (red), and EGFP-labeled T cells (green). Scale bar = 250µm. (D) Representative epifluorescence images illustrating three distinct tumor immunotypes: T cell-inflamed, DC-Innflamed; T cell-inflamed, DC-excluded; and T cell-excluded, DC-excluded. Images show KPP-mTagBFP2+ tumor cells (blue), EGFP+ T cells (green), tdTomato+ cDC1s (red), and a merged image. Scale bar = 250 µm.

Tumor rejection in STAMP requires the presence of cDC1s, as Batf3-deficient mice fail to reject STAMP tumors (**Supplementary Fig. 1A**). To determine whether increasing DC abundance could enhance rejection, we treated WT mice with Flt3L-Fc, a hematopoietic growth factor that expands multiple DC populations. However, Flt3L-Fc treatment did not improve tumor rejection compared to untreated WT controls (**Supplementary Fig. 1B**). These findings highlight two key contrasts: (1) Batf3 KO versus WT untreated, demonstrating the necessity of cDC1s for tumor rejection, and (2) WT FLT3L-treated versus WT untreated, showing that simply increasing DC numbers is insufficient to enhance rejection. Thus, cDC1s appear to play a qualitative rather than a purely quantitative role in tumor rejection and therefore may not be inherently rate limiting in this context. We therefore hypothesized that the spatial organization of DCs within the tumor or stroma might impact their antigen capture, maturation, or interactions with T cells, ultimately contributing to tumor rejection or progression.

We developed a genetic model for multiplexed imaging of cDC1s, T cells, and tumor cells in STAMP microtumor arrays (**Fig. 1B–C, Supplementary Fig. 1C–D**). We utilized Xcr1-Cre; Rosa26-LSL-tdTomato mice, which specifically label cDC1s with tdTomato fluorescence, enabling direct visualization of cDC1 distribution within tumors. To ensure clear spectral separation between cDC1s, T cells, and tumor cells, we crossed these mice to Rag2 KO mice and adoptively transferred EGFP-labeled T cells, allowing unambiguous tracking of T cell infiltration and distribution. This also prevented spurious tdTomato labeling of CD4+ T cells, ensuring that all labeled T cells originated from adoptive transfer (Mattiuz et al Front Immunol 2018). Because cDC1s are relatively rare in tumors, we treated mice with Flt3L-Fc to increase their abundance for imaging and spatial analysis; as discussed above, this treatment did not on its own affect tumor rejection rates. mTagBFP2+ tumor cells were used to establish tumors, creating a three-color system (tdTomato for cDC1s, EGFP for T cells, and mTagBFP2 for tumors) that enabled high-resolution imaging of immune-tumor interactions *in vivo* and *in situ*.

Using this fluorescence reporter model, we simultaneously visualized cDC1s, T cells, and tumor cells in living tumors, enabling direct classification of immunotypes in real time. We applied our previously established computational segmentation, tracking, and radial analysis approach (Ortiz-Muñoz et al., Nature 2023) to classify tumors as inflamed or excluded for each cell type and then assigned a combined immunotype based on the spatial distribution of both T cells and cDC1s (e.g., TC-In/DC-In, TC-In/DC-Ex, TC-Ex/DC-Ex). We observed that cDC1 spatial patterning correlates with T cell spatial patterning (TC-In/DC-In and TC-Ex/DC-Ex) in the majority, or about 75% of tumors (**Fig. 1D,E**). In 25% of the tumors, however, T cells were found to infiltrate the tumor while cDC1s were excluded and restricted to the periphery (**Fig. 1D,E**). It was therefore of interest to determine whether the differential spatial distributions of DCs and T cells were associated with different functional outcomes.

### cDC1/T Cell Immunotypes Predict with Tumor Outcomes and T Cell Cytolytic Activity

We have previously shown that T cell-inflamed STAMP tumors were more likely to be rejected, but this correlation was only significant at early time points after tumor seeding (Ortiz-Muñoz et al., Nature 2023). At later time points, a subset of tumors remained T cell-inflamed yet failed to be rejected, suggesting that T cell infiltration alone is insufficient for tumor clearance. To determine whether cDC1 spatial organization influences this outcome, we analyzed tumor rejection rates across immunotypes using our longitudinal STAMP imaging dataset. For each tumor, we tracked growth dynamics and immune infiltration over time, classifying tumors into TC-In/DC-In, TC-In/DC-Ex, or TC-Ex/DC-Ex groups based on our previously defined criteria. Tumor rejection was defined as complete regression before the experimental endpoint, while persistence was defined as continued tumor growth. Immunotype classifications were assessed at early time points (one week after tumor implantation, corresponding to the onset of T cell recruitment) and late time points, defined as the final observation before tumor rejection or the experimental endpoint. Our analysis revealed that TC-In/DC-In tumors—characterized by co-infiltration of cDC1s and T cells—were significantly more likely to undergo rejection at both early and late time points. In contrast, TC-In/DC-Ex tumors, which contained intratumoral T cells but lacked cDC1 co-infiltration, failed to be rejected despite their inflamed phenotype (**Fig. 2A**). These findings underscore that the presence of both cDC1s and T cells within the tumor is critical for effective tumor control.

**Figure 2.**
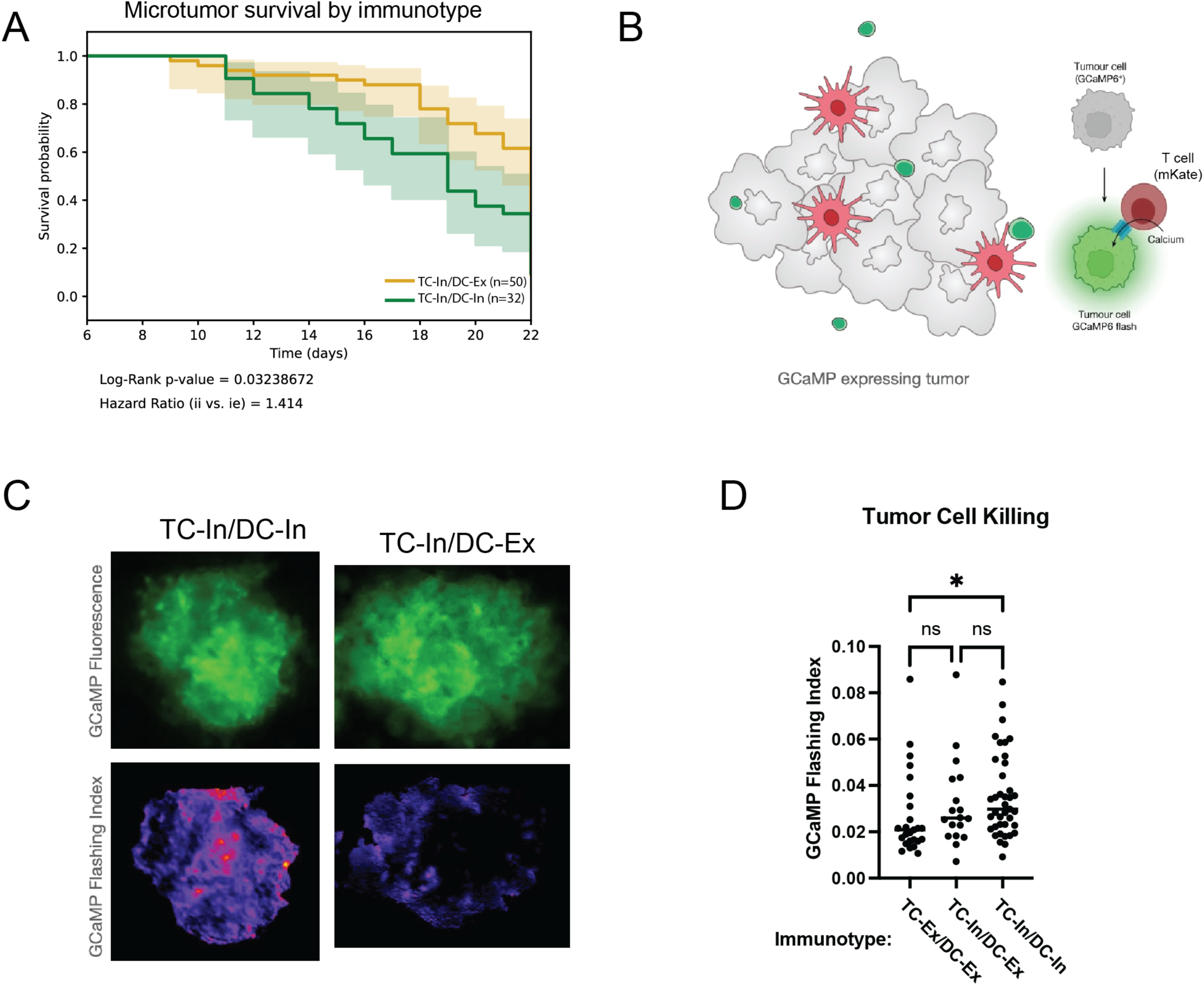
DC/T Cell Immunotypes Predict Tumor Rejection and Cytolytic Activity. (A) Kaplan-Meier curves rejection of individual microtumors classified by cDC1 and T cell immunotype in KPP-mTagBFP2 STAMP arrays implanted in Rag2-deficient Xcr1-cre Rosa-tdTomato animals with mKate T cells. *n* = 5 animals. (B) Schematic of the calcium-flashing reporter system for T cell cytolytic activity. Tumor cells express the GCaMP6 calcium sensor, which fluoresces upon calcium influx. Calcium influx serves as a proxy for T cell-induced pore formation, indicating active tumor cell killing. STAMP arrays containing GCaMP6+ KPP-mTagBFP2 tumor cells were implanted into Rag2-deficient Xcr1-Cre Rosa-tdTomato animals reconstituted with mKate+ T cells. (C) Representative images of calcium-flashing activity in T cell-inflamed, DC-Innflamed (TC-In/DC-In) and T cell-inflamed, DC-excluded (TC-In/DC-Ex) tumors. Top panels show GCaMP6 fluorescence (green), and bottom panels show standard deviation projections (higher standard deviation = more flashing). One representative tumor is shown for each group (n = 5 mice). Scale bar = 250 µm. (D) Quantification of the flashing index in TC-In/DC-In and TC-In/DC-Ex tumors. The flashing index reflects the frequency and intensity of calcium flashes. Data are mean ± SEM, n = 5 animals.

Next, we sought to determine whether the recruitment of T cells in inflamed tumors translated into increased tumor cell killing. Using the live-imaging capability of STAMP, we directly measured T cell cytolytic activity by employing a calcium-flashing reporter system. Tumor cells expressed GCaMP6, which fluoresces upon calcium influx triggered by T cell-induced pore formation, indicating individual attempted killing events (**Fig. 2B**). In TC-In/DC-In tumors, we observed frequent calcium flashes, while TC-In/DC-Ex tumors exhibited significantly fewer flashes (**Supplementary Video 1**). Quantification using a standard deviation projection revealed numerous hotspots of T cell activity in TC-In/DC-In tumors, compared to sparse activity in TC-In/DC-Ex lesions (**Fig. 2C**). Moreover, we found that TC-In/DC-In tumors with the highest flashing index exhibited the highest rejection, demonstrating that increased T cell activity is correlated with tumor control (**Fig. 2D**).

### Increased Clustering of CD4+ T Cells, CD8+ T Cells, and cDC1s in TC-In/DC-In Tumors

It is becoming evident that interactions between DCs and T cells in tumors, not just within draining lymph nodes, is an important feature of the anti-cancer immune response (Mellman et al., 2013). Indeed, this feature may reflect the the coordinated interactions between cDC1s, CD4+ T cells, and CD8+ T cells—referred to as immune triads—seen in both mouse and human tumors (Magen et al., Nat Med 2023; Espinosa-Carrasco et al., Cancer Cell 2024). Triads are proposed to reflect specialized immune niches, where cDC1s engage CD4+ T helper cells and CD8+ T cells to support activation, differentiation, and persistence. These studies have correlated immune triads with improved responses to checkpoint blockade, showing that they provide essential signals supporting CD8+ T cell cytotoxic potential and tumor elimination. However, these analyses have relied on fixed tumor samples, limiting the ability to determine if immune clusters actively contribute to tumor rejection.

By leveraging STAMP, we have been able to connect immune triads to specific immunotypes, which we can track dynamically by live imaging, allowing us to infer real-time changes in triad formation and function.

To assess immune clustering across different immunotypes, we first performed H&E staining to delineate microtumor boundaries (**Fig. 3A, upper**). We then conducted multiplex immunofluorescence (mIF) staining on STAMP tumor arrays using CD3 (pan-T cells), CD8 (cytotoxic T cells), and CLEC9A (cDC1s) to visualize immune cell distribution (**Fig. 3A, lower**). For classification, tumor boundaries were first annotated using H&E and DAPI staining. Immunotypes were then assigned based on CD3, CD8, and Clec9a fluorescence patterns followed by visual assessment to score whether immune cells were clustered or dispersed within the tumor microenvironment. Immune clusters composed of CD8+ T cells, CD4+ T cells, and Clec9a+ cDC1s were most abundant in TC-In/DC-In tumors, where they localized within the tumor core (**Fig. 3B**). In TC-In/DC-Ex tumors, T cells infiltrated but cDC1s were absent, leading to minimal clustering, while in TC-Ex/DC-Ex tumors, both cDC1s and T cells were restricted to the stroma (**Fig. 3C**). Quantification confirmed significantly reduced cDC1-T cell clustering in TC-In/DC-Ex and TC-Ex/DC-Ex tumors compared to TC-In/DC-In tumors (**Fig. 3D-F**).

**Figure 3.**
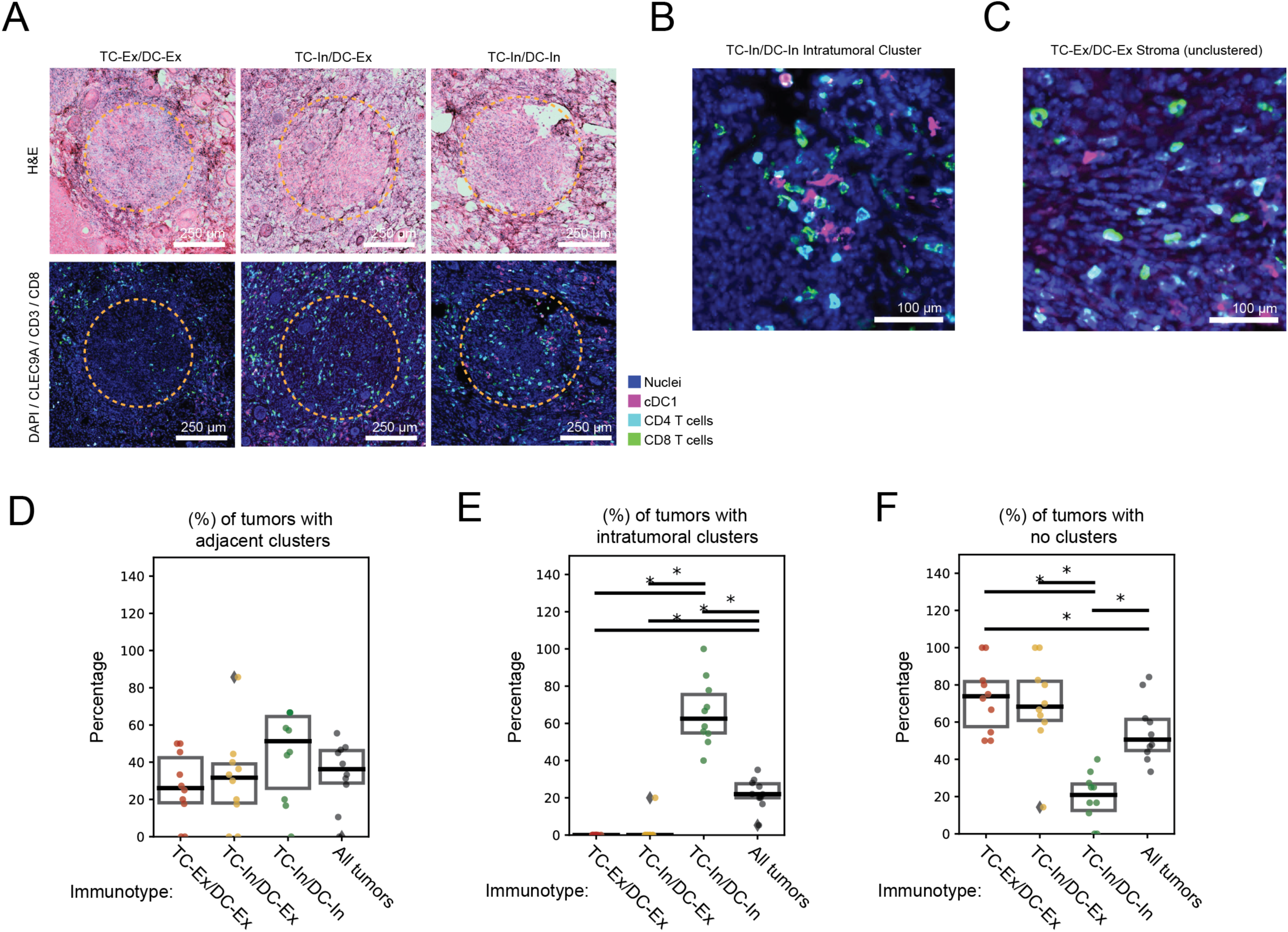
Increased Clustering of CD4+ T Cells, CD8+ T Cells, and cDC1s in TC-In/DC-In Tumors. (A) Representative images of three distinct STAMP tumor immunotypes: T cell-inflamed, DC-Innflamed (TC-In/DC-In); T cell-inflamed, DC-excluded (TC-In/DC-Ex); and T cell-excluded, DC-excluded (TC-Ex/DC-Ex). Top panels show multiplex immunofluorescence (IF) staining for DAPI (nuclei), Clec9a+ cDC1s (magenta), CD4+ T cells (cyan), and CD8+ T cells (green). Bottom panels show corresponding H&E staining of the same tumors. Tumor outlines are indicated by yellow dotted lines, n=10 mice. Scale bar = 250 µm. (B) Representative inset from an TC-In/DC-In tumor, showing close spatial interactions between CD4+ T cells, CD8+ T cells, and Clec9a+ cDC1s, n=10 mice. Scale bar = 50 µm. (C) Representative image of an TC-Ex/DC-Ex tumor stroma, showing CD4+ T cells, CD8+ T cells, and Clec9a+ cDC1s in a non-clustered distribution, n=10 mice. Scale bar = 250 µm.

These findings establish immune clustering as a defining feature of TC-In/DC-In tumors, providing a microscale interaction framework for interpreting the functional differences in T cell activation and tumor rejection described in later sections. By linking immune clustering to real-time functional outcomes in individual tumors, the live cell approach has provided direct evidence that cDC1-T cell interactions shape tumor immunity beyond correlative observations in fixed tissues.

### cDC1 Phenotypes Reflect Specialized Functions in Distinct Tumor Immunotypes

To investigate the transcriptional and phenotypic diversity of cDC1s in tumors with different immune infiltration patterns, we performed flow cytometry, bulk RNA sequencing, and single-cell RNA sequencing (scRNA-seq) on STAMP tumors classified by both T cell and cDC1 infiltration patterns. For bulk transcriptional profiling, tumor biopsies were sorted into three populations (cDC1s, T cells, and all live cells) to enable a comprehensive analysis of immunotype-specific gene expression. We also performed scRNA-seq on biopsied tumors, however the number of recovered cells was insufficient to compare transcriptional profiles across immunotypes. As a result, bulk RNA sequencing provided insights into immunotype-specific cDC1 transcriptional programs, while scRNA-seq was used to broadly characterize the immune composition of STAMP tumors and assess the transcriptional states of the major immune populations.

RNA sequencing revealed that Xcr1+ cDC1s in TC-In/DC-In tumors exhibited gene expression patterns suggestive of specialized functions in T cell activation (Cd86, Icosl) and T cell recruitment (Ccl5) (**Fig. 4A, Supplementary Fig. 2A-D**). These cells show upregulation of stress-response genes (Hspa5, Atf4, Atf6), survival regulators (Bcl3, Cish), and signaling modulators (PP2A subunits). This transcriptional program suggests that tumor-infiltrating cDC1s in TC-In/DC-In tumors maintain a mature, costimulatory phenotype while resisting stress-induced dysfunction or death, enabling them to sustain productive interactions with T cells. (**Fig. 4A, Supplementary Fig. 2A-D**). On the other hand, in TC-In/DC-Ex tumors, cDC1s exhibited gene expression patterns associated with mature (Cd274, Relb) and migratory cells optimized for lymph node trafficking (Ccr7, Fscn1), cross-presentation (B2m, Tap2, Wdfy4), and CD4+ T cell activation (Cd40, H2-Ab1, Ciita). These patterns also indicate motility (Actb, Itgb1, Arpc2, Cdc42) but suggest increased susceptibility to cell death (Casp8, Ripk3) (**Fig. 4B, Supplementary Fig. 2A-D**). In TC-Ex/DC-Ex tumors, cDC1s display gene expression patterns consistent with a stressed (Sod2, Nfe2l2, Atf6, Hspa5), inflammation-adapted state (Myd88, Il1b, Tgfb1, Stat1) that is less mature (lower expression of Cd86, Ccr7, Cd274, and Cd40), and possibly more adept at antigen capture (Fcgr2b, Tyrobp) and endosomal trafficking (Lamp1, Eea1) (**Fig. 4C, Supplementary Fig. 2A-D**).

**Figure 4.**
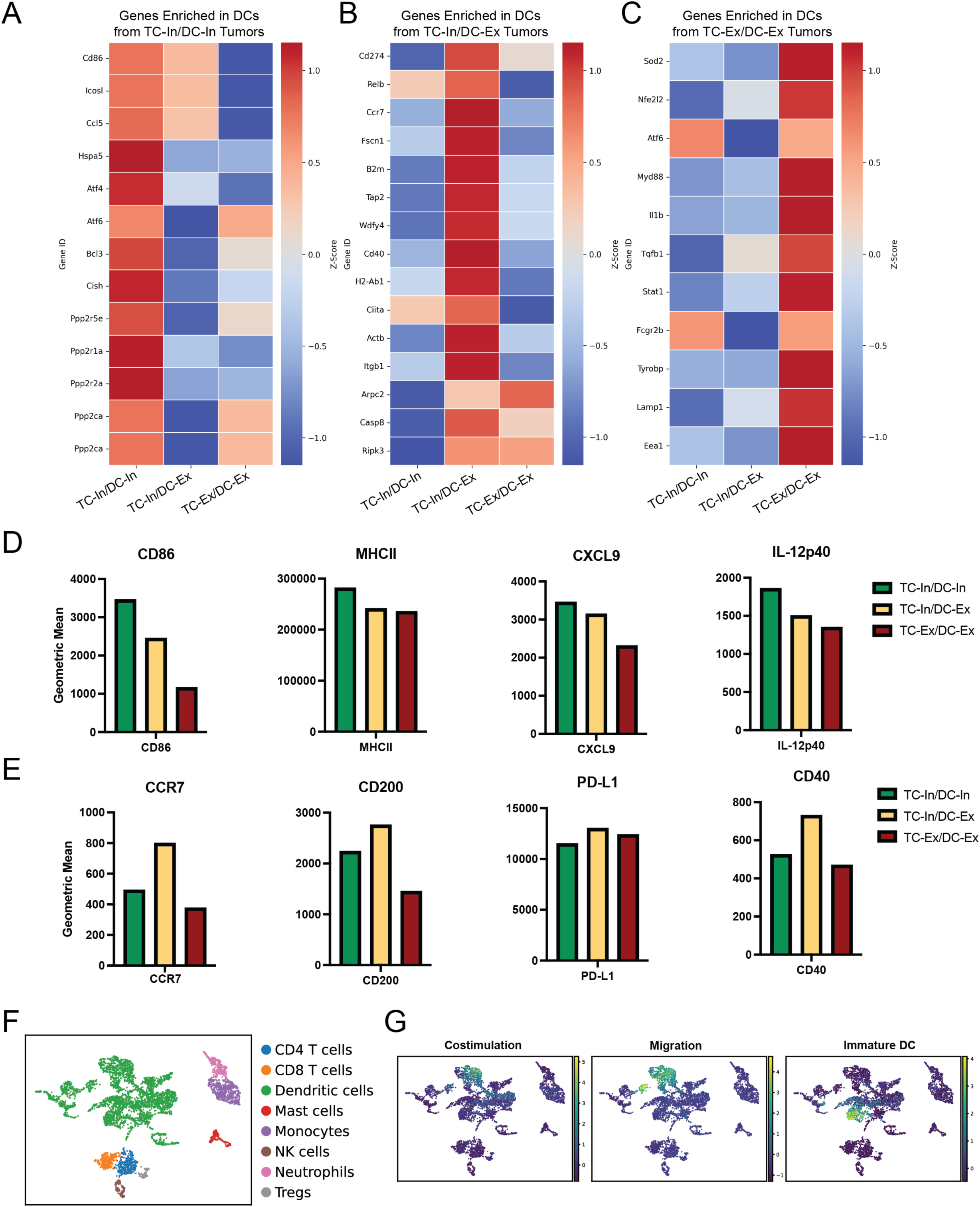
Dendritic Cell Phenotypes Reflect Specialized Functions in Distinct Tumor Immunotypes. (A–C) Bulk RNA sequencing of Xcr1+ cDC1s isolated from STAMP microtumors biopsied from KPP-mTagBFP2 tumor arrays implanted in Xcr1-Cre; Rosa26-LSL-tdTomato Rag-deficient mice reconstituted with GFP+ T cells. Tumors were classified by immunotype: T cell-inflamed, DC-Innflamed (TC-In/DC-In); T cell-inflamed, DC-excluded (TC-In/DC-Ex); and T cell-excluded, DC-excluded (TC-Ex/DC-Ex). Heatmaps show genes upregulated in Xcr1+ cDC1s from (A) TC-In/DC-In tumors, (B) TC-In/DC-Ex tumors, and (C) TC-Ex/DC-Ex tumors. Data represent pooled tumors from 5 mice, with three replicates. (D–E) Flow cytometry analysis of pooled tumors from KPP-mTagBFP2 tumor arrays implanted in Xcr1-Cre; Rosa26-LSL-YFP Rag-deficient mice reconstituted with mKate+ T cells. Bar graphs show the geometric mean fluorescence intensity (gMFI) of (D) TC-In/DC-In associated markers MHCII, CD86, IL-12, and CXCL9 in Xcr1+ cDC1s from TC-In/DC-In, TC-In/DC-Ex, and TC-Ex/DC-Ex tumors. (E) Bar graphs show gMFI of TC-In/DC-Ex associated markers CCR7, CD200, PD-L1 in Xcr1+ cDC1s across immunotypes. Data represent pooled tumors from 15 mice; one representative experiment of two independent replicates is shown. (F) UMAP projection of single-cell RNA sequencing data from CD45+ immune infiltrates of STAMP tumors, with T cells and cDC1s overrepresented due to targeted enrichment for a cDC1 fraction (CD45+CD11c+MHCII+) and a T cell fraction (CD45+CD90+), which were combined for downstream analysis. Data represent all tumors pooled from n=10 mice. (G) Feature plots displaying the expression of gene signatures associated with costimulation (Cd86, Icosl), migration (Ccr7, Fscn1, Cd200), and immature cDC1 lineage markers (Flt3, Irf8).

Flow cytometry analysis of tumors pooled by immunotype from 15 STAMP array-bearing mice corroborated these findings and further highlighted trends across the three immunotypes. Canonical maturation markers such as cell surface MHCII and CD86 were highest in TC-In/DC-In tumors, intermediate in TC-In/DC-Ex tumors, and lowest in TC-Ex/DC-Ex tumors (**Fig. 4D**). Similarly, cDC1 secretory products that support T cell function (e.g., IL-12 and CXCL9) were highest in TC-In/DC-In tumors. Migratory cDC1-associated markers such as CCR7, CD200, PD-L1 and the DC licensing receptor CD40 were predominantly upregulated in TC-In/DC-Ex tumors, where cDC1s appeared to adopt a more migratory phenotype (**Fig. 4E**). These markers were lower in TC-In/DC-In tumors, suggesting cDC1 retention and interaction with T cells within the tumor (**Fig. 4E**). Flow cytometry also validated the TC-Ex/DC-Ex cDC1s exhibit a less mature phenotype with low expression of both sets of maturation markers.

We performed single-cell RNA sequencing of the immune infiltrate in biopsied STAMP tumors (**Fig. 4F**, **Supplementary Fig. 2E**). UMAP analysis revealed non-overlapping expression of costimulatory, migration-associated, and immature dendritic cell gene signatures across cDC1s (**Fig. 4G**). These features supported the presence of transcriptionally defined clusters with enriched expression of costimulatory, migration-associated, or immature dendritic cell gene signatures. Thus, cDC1s in different tumor immunotypes display distinct functional states rather than representing a uniform population across all immunotypes. Whether these states reflect stable differentiation programs or dynamic transitions remains to be determined.

Taken together, the data indicate that cDC1 function in STAMP tumors is highly context-dependent, with costimulatory cDC1s more prevalent in intratumoral environments characterized by direct cDC1/T cell interactions, as seen in TC-In/DC-In tumors. Conversely, migratory cDC1s predominate at the periphery of tumors where T cells infiltrate without cDC1 co-infiltration (TC-In/DC-Ex). Thus, depending on their spatial distribution within and around the tumor, cDC1s can exhibit markedly different functional attributes.

### DC-In Tumors Contains Cytotoxic Effectors, While DC-Ex Tumors Are CD4-Dominated with Mixed Effector and Regulatory Profiles

We next asked if the functional attributes of the T cells in TC-In/DC-In tumors were also distinct from T cells in TC-in/DC-ex tumors. For this purpose, we analyzed the bulk transcriptional profiles of T cells isolated from TC-In/DC-In, TC-In/DC-Ex, and TC-Ex/DC-Ex immunotypes. RNA sequencing revealed that T cells in TC-In/DC-In tumors exhibited gene expression patterns associated with cytotoxic effector function (e.g., Cd8a, Cd8b1, Cd247, Ccl5), TCR signaling (Zap70, Lat, Lck, Cd226), and early activation (Tbx21, Pdcd1, Slamf6). We also detected expression of Cx3cr1, a chemokine receptor implicated in tissue infiltration and retention of T cells (**Fig. 5A**, **Supplementary Fig. 3A-D**).

**Figure 5.**
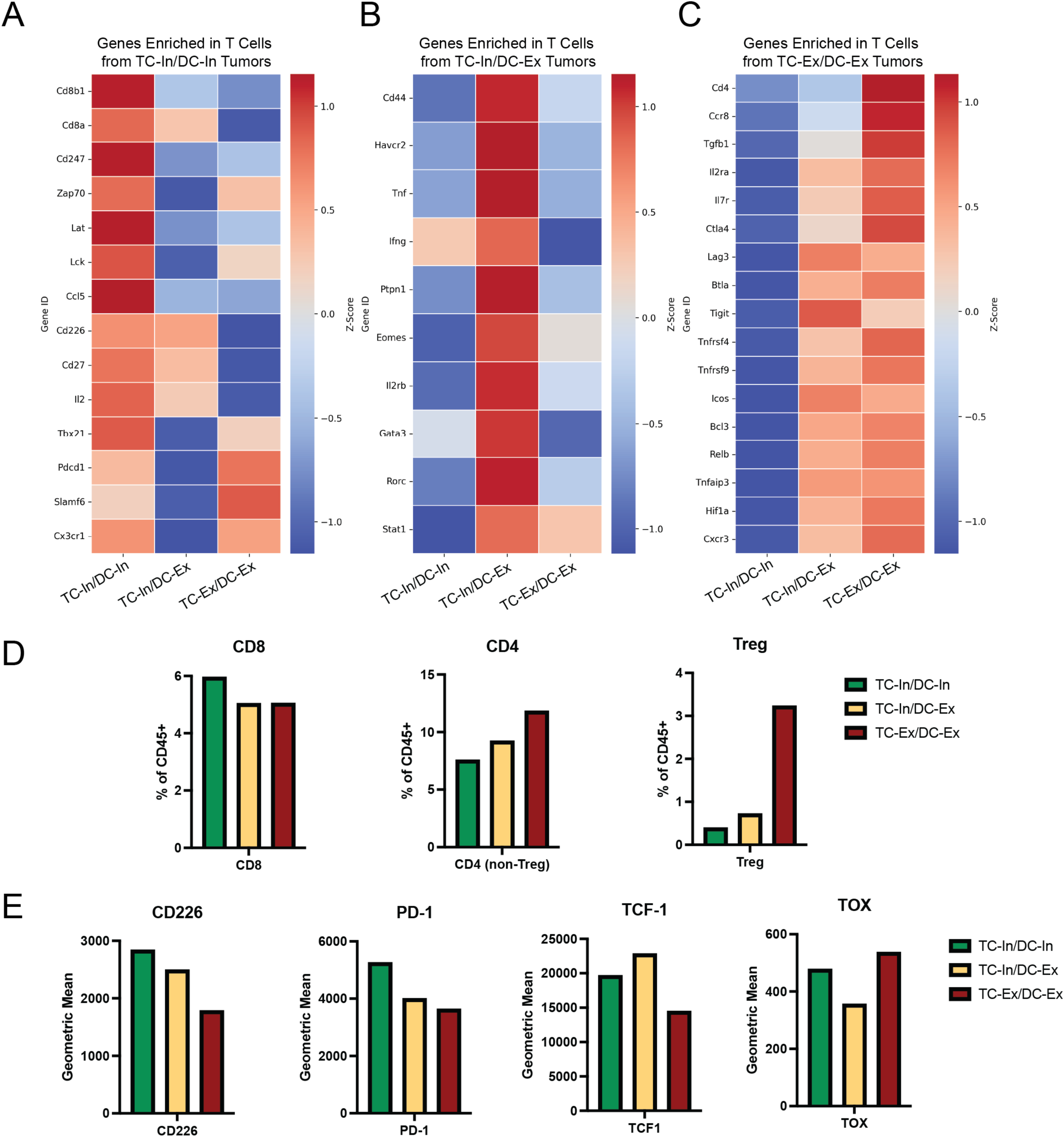
cDC1 Infiltration Correlates with Cytotoxic Effector T Cells in TC-In/DC-In Tumors, While Excluded Tumors Are CD4-Dominated with Mixed Effector and Regulatory Profiles. (A–C) Bulk RNA sequencing of T cells isolated from STAMP microtumors biopsied from KPP-mTagBFP2 tumor arrays implanted in Xcr1-Cre; Rosa26-LSL-tdTomato Rag-deficient mice reconstituted with GFP+ T cells. Tumors were classified by immunotype: T cell-inflamed, DC-Innflamed (TC-In/DC-In); T cell-inflamed, DC-excluded (TC-In/DC-Ex); and T cell-excluded, DC-excluded (TC-Ex/DC-Ex). Heatmaps show genes upregulated in T cells from (A) TC-In/DC-In tumors, (B) TC-In/DC-Ex tumors, and (C) TC-Ex/DC-Ex tumors. Data represent pooled tumors from 5 mice, with three replicates. (D–E) Flow cytometry analysis of pooled tumors from KPP-mTagBFP2 tumor arrays implanted in Xcr1-Cre; Rosa26-LSL-YFP Rag-deficient mice reconstituted with mKate+ T cells. (D) Bar graphs show abundance of CD8 T cells, CD4 T cells and Tregs as a proportion of CD45+ cells. (E) Bar graphs show gMFI of PD-1, CD226, TCF-1 and TOX in CD3+ T cells across immunotypes. Data represent pooled tumors from 15 mice; one representative experiment of two independent replicates is shown.

In contrast, T cells from TC-In/DC-Ex tumors showed increased expression of genes typically associated with CD4+ T cell effector function (Cd44, Tnf, Ifng, Stat1), and transcriptional features suggestive of a CD8+ T cell activated or hyperactivated state, including Eomes, Havcr2 (TIM-3), and Ptpn1. Expression of lineage-defining genes including Il2rb, Rorc, and Gata3 suggests that multiple distinct CD4+ T cell subsets are present within these tumors (**Fig. 5B, Supplementary Fig. 3A-D**). The elevated expression of inhibitory receptors such as Lag3, Ctla4, and Tigit suggests that T cells in TC-In/DC-Ex tumors may be progressing along an exhaustion trajectory (in the case of CD8+ T cells) or adopting immunoregulatory features (in the case of CD4+ T cells) (**Fig. 5C, Supplementary Fig. 3A-D**).

In TC-Ex/DC-Ex tumors, T cells exhibited gene expression profiles suggestive of a predominantly CD4+ population. These included elevated expression of canonical regulatory T cell (Treg) and Type 1 regulatory (Tr1)-associated genes (Ccr8, Tgfb1, Il2ra) as well as inhibitory co-receptors (Ctla4, Lag3, Tigit). We also observed enrichment of genes linked to chronic activation (Btla, Tnfrsf4/OX40, Tnfrsf9/4-1BB, Bcl3, Relb, Tnfaip3) and metabolic adaptation (Hif1a). Elevated Cxcr3 expression may contribute to stromal retention, further limiting T cell access to the tumor core (**Fig. 5C**). While many of these features are consistent with regulatory or exhausted CD4+ T cells, they may also reflect terminal differentiation or dysfunction among CD8+ T cells. Because the RNA-seq was performed on bulk-sorted T cells, both interpretations remain possible, but the data do indicate significant differences in T cell state as a function of tumor immunotype.

Flow cytometry confirmed that both CD4+ and CD8+ T cells were present across all three immunotypes, with a modest relative enrichment of CD8+ T cells in TC-In/DC-In tumors and a clear increase in Foxp3+ regulatory T cells in TC-Ex/DC-Ex tumors, consistent with the gene expression data (**Fig. 5D**). Phenotypic profiling revealed trends in activation and differentiation marker expression (**Fig. 5E**): CD226 and PD-1, markers associated with activated cytotoxic T cells, were most highly expressed in TC-In/DC-In tumors, consistent with their role in effective anti-tumor responses (Weulersse et al., 2020; Banta et al., 2022). TCF-1, a marker of progenitor-like or precursor exhausted T cells (Tscm or Tpex), was enriched in TC-In/DC-Ex tumors, suggesting that cDC1 exclusion may preserve a less differentiated T cell pool. In contrast, TOX, a transcription factor associated with terminal exhaustion, was most highly expressed in TC-Ex/DC-Ex tumors, with intermediate expression in TC-In/DC-In and the lowest levels in TC-In/DC-Ex tumors.

While the phenotypic differences observed were generally modest, they are consistent with the RNA-seq findings and support a model in which TC-In/DC-In tumors promote effector T cell differentiation, TC-In/DC-Ex tumors maintain a less differentiated progenitor-like pool, and TC-Ex/DC-Ex tumors are skewed toward terminally exhausted or regulatory phenotypes.

### Antigen-Matched T Cells Require cDC1s for Efficient Recruitment and Activation

To investigate how cDC1s influence T cell recruitment and effector function independently of priming, we devised an approach in which T cells were already activated prior to tumor challenge. We engineered the KPP tumor cell line to express a defined tumor neoantigen (M86; Capietto et al., 2020) and implanted these cells into STAMP arrays. In parallel, donor mice were vaccinated with RNA-lipoplexes encoding the M86 antigen to generate a pool of antigen-specific, primed T cells. These T cells were isolated and adoptively transferred into tumor-bearing recipient mice. As controls, we also transferred T cells from OVA-vaccinated mice (antigen-mismatched) or naïve mice (unprimed).

We compared tumor rejection outcomes in Rag2 KO mice, which lack endogenous T and B cells but retain cDC1s, and BatF3 KO mice, which lack cDC1s but retain endogenous T cells. Despite receiving identical T cell transfers, antigen-matched T cells mediated significantly more effective tumor rejection in Rag2 KO mice than in BatF3 KO mice (**Supplementary Fig. 4A–B**; HR 62.63 vs. 7.64, M86 vs. naïve), indicating that cDC1s enhance T cell-mediated tumor clearance even after priming. While the improved efficacy in Rag2 KO mice may partially reflect the absence of endogenous T cells competing for niche space, the differential outcomes strongly implicate cDC1s as important facilitators of anti-tumor immunity, independent of their role in T cell priming.

We also observed that vaccine-elicited T cells—both antigen-matched and mismatched—were more effective at infiltrating tumors than naïve T cells (**Supplementary Fig. 4C**), suggesting a general, antigen-independent effect of prior activation on trafficking. Importantly, T cell infiltration was significantly greater in Rag2 KO mice than in BatF3 KO mice (Supplementary Fig. 4D–E), reinforcing the role of cDC1s in facilitating intratumoral T cell recruitment. While cDC1s were not strictly required for T cell trafficking, their absence impaired both T cell infiltration and tumor rejection.

Together, these findings provide direct and functional evidence suggesting that cDC1s, beyond their role in T cell priming, contribute to anti-tumor immunity by facilitating intratumoral recruitment and sustaining effector function of antigen-matched T cells within the tumor microenvironment.

### Engineered Chemokine Expression Enhances Tumor Rejection, but Effective Outcomes Require cDC1 and T Cell Co-Recruitment

Given the likely role of cDC1s in recruiting T cells to tumors, we investigated how the chemokine environment of individual lesions in the STAMP array may contribute to immune cell infiltration and tumor rejection. We focused on CCL5, XCL1, and CXCL9 due to their well-established functions in attracting cDC1s and T cells (Böttcher et al., Cell 2018; Gorbachev et al., 2007; Chow et al., Immunity 2019). To manipulate chemokine expression in a controlled and physiologically relevant manner, we developed a doxycycline-inducible system for tumor-specific chemokine expression. This approach enabled us to restrict chemokine production to the in vivo setting, minimizing potential autocrine effects during in vitro culture. This is a particular concern for CCL5 and CXCL9, which have been shown to influence tumor cell proliferation or migration (Walser et al., Cancer Res. 2006, Karnoub et al., Nature 2007). Doxycycline treatment was initiated one day before tumor implantation to ensure uniform induction of chemokine expression. To enable direct comparison of immune responses within the same host, we implanted chemokine-expressing tumor cells in the left ear and control cells in the right ear of the same mouse (**Fig. 6A**).

**Figure 6.**
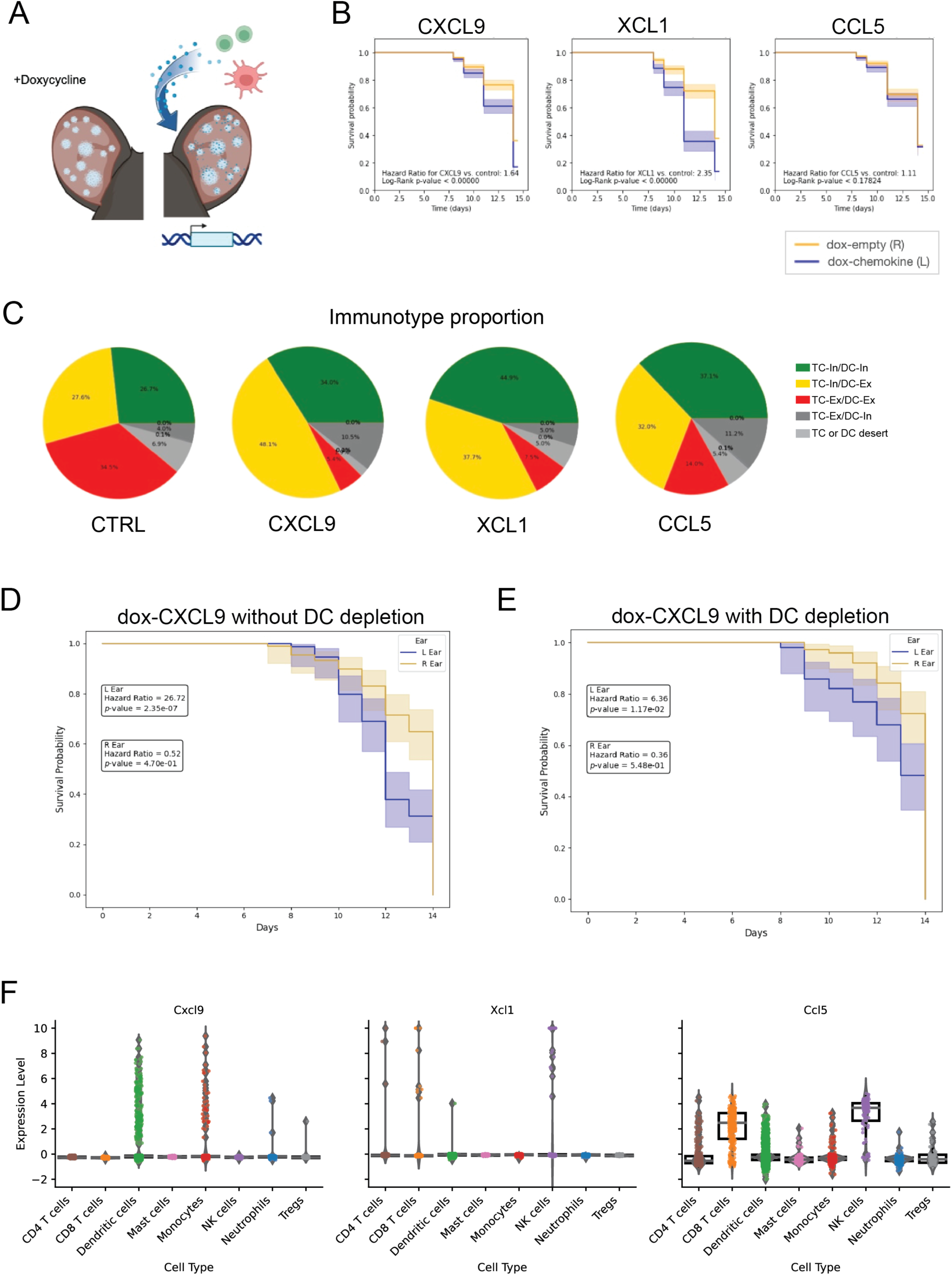
Engineered Chemokine Expression Enhances Tumor Rejection, but Effective Outcomes Require cDC1 and T Cell Co-Recruitment. (A) Chemokine gene expression from single-cell RNA sequencing of STAMP tumors classified by immunotype: T cell-inflamed, DC-Innflamed (TC-In/DC-In); T cell-inflamed, DC-excluded (TC-In/DC-Ex); and T cell-excluded, DC-excluded. (TC-Ex/DC-Ex). Data show immunotype-specific enrichment of CCL5, XCL1, and CXCL9. Data represent pooled tumors from 10 mice (B) Schematic of the doxycycline (dox)-inducible system for tumor-specific chemokine expression. (C) Kaplan-Meier survival curves comparing tumor rejection rates in contralateral ears of wild-type mice implanted with dox-inducible KPP-mTagBFP tumor arrays expressing CXCL9, XCL1, or CCL5 versus empty vector control. n = 5 mice per group. (D) Proportions of tumor immunotypes in Rag-deficient Xcr1-Cre; Rosa26-LSL-tdTomato mice reconstituted with EGFP+ T cells. KPP-mTagBFP tumor arrays expressing CXCL9, XCL1, CCL5, or empty vector control were induced with doxycycline. Immunotypes were determined based on cDC1 and T cell infiltration patterns. n = 5 mice per group. (E–F) Kaplan-Meier survival curves comparing tumor rejection rates in Clec9a-Cre; Zbtb46-DTR mice implanted with dox-inducible KPP-mTagBFP tumors expressing CXCL9 versus empty vector control. (E) No diphtheria toxin mediated dendritic cell depletion; (F) dendritic cell depletion initiated on day 4 post-implantation using diphtheria toxin. n = 5 mice per group.

In wild-type mice, tumor cell expression of the T cell chemoattractant CXCL9 significantly enhanced tumor rejection, with a modestly higher survival benefit compared to the contralateral tumors lacking chemokine induction (H.R. 1.64, **Fig. 6B)**. XCL1, a key chemoattractant for cDC1s, had an even greater effect, inducing robust rejection in the majority of tumors (H.R. 2.35, **Fig. 6B**). However, induction of CCL5 did not significantly enhance rejection, perhaps owing to its broader spectrum of target cells and lack of selectivity for DCs or T cells (H.R. not significant, **Fig. 6B**).

To understand how these outcomes related to immune infiltration patterns, we repeated the experiment in Rag2-deficient mice reconstituted with tdTomato-expressing polyclonal T cells from naïve donors. Immunotype analysis revealed that CXCL9 expression primarily increased the frequency of TC-In/DC-Ex tumors, XCL1 promoted a shift toward TC-In/DC-In tumors, and CCL5 increased the proportion of both TC-In/DC-Ex and TC-In/DC-In tumors (**Fig. 6C**). Together, these results show that while CXCL9 enhances T cell infiltration, its effects on tumor rejection are approximately 30% lower than those of XCL1, which facilitates co-infiltration of cDC1s and T cells (H.R. 1.64 vs. 2.35; Fig. 6B). Interestingly, although CCL5 shifted immunotypes toward more inflamed patterns, this did not improve tumor rejection, again, possibly reflecting its more pleiotropic effects on immune cell recruitment or autocrine effects on the tumor cells themselves.

Finally, we evaluated the role of cDC1s in CXCL9-driven tumor rejection using an inducible cDC1 depletion model. We employed a conditional system where Clec9a-Cre drives Zbtb46-DTR expression, enabling DT-mediated cDC elimination. The dual specificity of Clec9a and Zbtb46 ensures precise targeting of cDCs while minimizing off-target effects; however, both cDC1 and cDC2 are known to be targeted by this Cre (Mattiuz et al 2018). To preserve initial T cell priming and recruitment, DT treatment was initiated on day 4 post-tumor implantation, allowing us to specifically assess the role of cDC1s in sustaining T cell infiltration and spatial interactions within CXCL9-expressing tumors.

In tumors expressing CXCL9, cDC1 depletion significantly diminished, but did not fully abrogate, the tumor survival benefit typically observed with CXCL9 induction (**Fig. 6D-E**). This indicated that while CXCL9 enhances T cell infiltration, its full impact on tumor rejection requires cDC1s. CXCL9 is well established as a key mediator of CXCR3+ T cell recruitment; however, its role may extend beyond infiltration, as intratumoral CXCL9 production by cDC1s has been shown to promote productive cDC1-T cell interactions necessary for effective anti-tumor immunity (Chow et al., Immunity 2019).

It is also possible, however, that ongoing cDC1 function in the tumor-draining lymph node (dLN), rather than in the tumor itself, is required for sustained T cell priming and effector function (Roberts et al., Cancer Cell 2016). Our findings show that while tumor-derived CXCL9 broadly facilitates T cell recruitment, the presence of cDC1s remains essential for optimal tumor rejection. Specifically, cDC1-produced CXCL9 may uniquely promote microscale interactions that enhance T cell activation and effector function (Meiser et al 2023). Thus, beyond recruitment, cDC1s may shape the local immune landscape within the tumor or sustain T cell responses through continuous priming in the dLN, both of which could contribute to immune triad formation and amplified anti-tumor immunity.

To determine the cellular sources of CXCL9, XCL1, and CCL5 within the tumor microenvironment, we analyzed their expression across immune populations using our single-cell sequencing atlas. CXCL9 was primarily produced by DCs and monocytes, suggesting that CXCL9 availability was largely dictated by the presence and activation of myeloid cells (**Fig. 6F**). This finding also supports the hypothesis that cDC1s may contribute through local costimulatory interactions in tumors and also through CXCL9-mediated mechanisms of T cell recruitment and activation. Consistent with previous reports (Böttcher et al., 2018), we found that in our system XCL1 was predominantly expressed by NK cells, with lower levels in CD4⁺ and CD8⁺ T cells, while CCL5 was mainly produced by NK cells and CD8⁺ T cells (**Fig. 6F**). These data highlight the distinct cellular sources of key chemokines and further support the notion that successful immune infiltration and tumor rejection require coordinated cDC1-T cell functions rather than T cell recruitment alone (**Fig. 6F**).

Our findings demonstrate that chemokine expression is crucial for shaping rejection-associated immunotypes by driving the spatial distribution of cDC1s and T cells within the TME. While ectopic chemokine expression from tumor cells effectively enhanced tumor rejection, the enhance efficacy reflected more than just the influx of T cells. For instance, the impact of CXCL9 was significantly dependent on the presence of cDC1s, which presumably act not only to attract T cells but also sustain their activity by the cross presentation of tumor antigens and provision of costimulatory ligands and immunostimulatory cytokines.

## Discussion

This study demonstrates the critical role of cDC1s and chemokines in orchestrating anti-tumor immunity through the recruitment, activation, and spatial organization of T cells within the TME. By allowing us to observe the relationship between spatial distribution and tumor rejection in individual tumors, STAMP has provided the unique opportunity to obtain functional evidence attesting to the significance of the interaction patterns between cDC1s and T cells. Our data suggest that distinct immunotypes that predict tumor outcomes, with TC-In/DC-In (doubly inflamed) tumors exhibiting the highest rejection rates. These findings highlight that intratumoral DC/T cell clustering is a hallmark of productive anti-tumor immunity. By underscoring the importance of cellular co-localization and cross-talk in driving immune responses in STAMP, the model supports likely functional significance of DC/T cell clustering observed in human cancer (Magen & Hamon, 2023; Chen et al., 2024).

DCs are specialized for bridging the innate and adaptive arms of the immune response (Mellman and Steinman, 2001). In the case of shaping anti-tumor immunity, cDC1s in particular act to capture tumor antigens and mature in response to inflammatory stimuli, migrate to dLN where they process and cross-present tumor antigens to T cells thus priming the adaptive response, and also present antigen in the tumor to both attract T cell influx and sustain T cell activity. Our data reveal that these functions are variable across tumor immunotypes. In TC-In/DC-In tumors, cDC1s support tumor-resident adaptive functions, characterized by enhanced DC maturation, facilitating the co-infiltration and interaction of cDC1s and T cells, and enhanced T cell effector function. This is evidenced by increased expression of maturation markers (e.g., MHCII, CD86), non-canonical co-stimulatory molecules (e.g., CD80, PVR), and chemokines such as CXCL9 and IL-12. These cDC1s also exhibit lower expression of migration-associated genes, suggesting retention within the tumor and sustained interactions with T cells, which are critical for driving effective anti-tumor immunity. T cells in this immunotype also appeared in our live imaging and flow cytometry experiments to be more potent effectors.

Conversely, in tumors inflamed by T cells but excluded for cDC1 (TC-In/DC-Ex), cDC1s predominantly display classical adaptive phenotypes associated with antigen trafficking to lymph nodes. These cDC1s exhibit a migratory maturation program with increased expression of genes such as Ccr7 and Fscn1, but they show less expression of genes involved in antigen presentation, co-stimulation, and effector cytokine production compared to TC-In/DC-In tumors. This separation of cDC1 functions may explain the reduced efficacy of T cells in TC-In/DC-Ex tumors, despite their substantial infiltration into the tumor. In addition, it is likely significant that the absence of cDC1’s from the tumor parechyma deprives T cells from local stimulation that may be needed for optimal anti-tumor activity.

Through adoptive transfer experiments, we confirmed that cDC1s are indispensable for the recruitment and activation of antigen-matched T cells. In Rag2-deficient mice, the presence of cDC1s correlated with robust T cell activation and tumor rejection, while BatF3-deficient mice lacking cDC1s exhibited impaired T cell recruitment and activation. These results highlight the role of cDC1s, not only in priming T cells in lymph nodes, but in maintaining their effector function within the tumor. Interestingly, while vaccine-elicited antigen-matched T cells were able to reach tumors in the absence of cDC1s, their numbers and activation levels were significantly reduced, underscoring the central role of cDC1/T cell interactions in sustaining anti-tumor immunity. This aligns with previous studies demonstrating that tumor-resident cDC1s are required to support effector T cell function and intratumoral accumulation (Hildner et al. 2008, Roberts et al 2016, Salmon et al 2016) even in models where T cell priming is bypassed and and dendritic cells act solely to support effector T cell function (Broz et al 2014, Spranger et al., 2017). Our study builds on this foundation by directly visualizing dynamic cDC1–T cell interactions during tumor cell killing in vivo and revealing a functional connection to their spatial organization in intratumoral immune clusters, structures not previously described in these earlier studies.

To better understand how cDC1–T cell clusters are coordinated within tumors, we turned to inducible chemokine expression to manipulate the local immune landscape. These experiments revealed that while both CXCL9 and XCL1 can enhance tumor rejection, they do so in distinct ways—XCL1 preferentially recruits cDC1s and fosters TC-In/DC-In immunotypes, whereas CXCL9 alone primarily draws in T cells, often in the absence of intratumoral cDC1s. Interestingly, even though both chemokines contribute to rejection, XCL1 had a more pronounced effect, suggesting that effective anti-tumor immunity may hinge more critically on bringing in dendritic cells than T cells alone (Böttcher et al 2018). The modest impact of CCL5 further underscores that not all chemokines are equally suited to reprogram the tumor microenvironment. Perhaps most tellingly, CXCL9-driven rejection was abrogated in the absence of cDC1s, reinforcing the idea that cDC1s play essential roles beyond recruitment— serving as local hubs for antigen presentation and co-stimulation. These findings point to an interdependence between T cell and cDC1 infiltration that shapes immune-responsive tumors.

Together, our work has significant implications for the design of immunotherapies targeting the chemokine-cDC1-T cell axis. While ectopic chemokine expression from tumor cells effectively enhances immune infiltration and tumor rejection, this strategy may not address critical elements of natural tumor immunity, such as the functional maturation and strategic localization of cDC1s within the tumor parenchyma. Therapies that enhance cDC1 recruitment and retention within tumors, while also supporting their activation and interaction with T cells, are likely to be more effective in achieving durable tumor rejection. These considerations may be of particular importance to efforts aimed at using adoptive T cell therapy in solid tumors.

### Limitations of the Study

While the STAMP model provides a powerful platform for dissecting localized immune interactions, it does not fully capture the complexity of human tumors, particularly in metastatic settings or those with heterogeneous immune infiltration. Additionally, while we identify cDC1-T cell clustering as a key feature of tumor rejection, its relevance in human tumors remains to be fully explored. Our use of transgenic models and doxycycline-inducible chemokine systems provides mechanistic insights but does not directly translate to therapeutic strategies, highlighting the need for approaches that can effectively implement chemokine-driven immune coordination in clinical settings. Future studies incorporating systemic tumor models and interventional strategies will be essential to bridge these findings to immunotherapy applications.

## Resource Availability

### Lead Contact

Further information and requests for resources and reagents should be directed to and will be fulfilled by the lead contact, Christine Moussion.

## Materials Availability

## Acknowledgements

We thank the members of the Moussion and Mellman laboratories for their advice, discussions, and reagents. We are grateful to Alan Gutierrez and Yajun Chestnut for assistance with takedowns and tissue harvest. We acknowledge the Genentech mouse husbandry team, including Fermin Gallardo-Chang, Shannon Powell, Tina Scholl, Bridget Halpenny, Mariela Pantle, Ben Torres, Michael Long, Meredith Dempsey, Chris De La Cruz, and Raymond Asuncion. We thank the veterinary care team, including Raul Garcia-Gonzalez, Libette Roman, Emmanuel Chua, Millicent Lu, and Jill Yamada, for their dedicated animal care. We also appreciate the support of the Genentech Laboratory Animal Resources team, including Bridget Maguire-Colton, Caroline Sepulveda, Dorie Montoya, Ryan Scott, Mike Reich, Suzette Weber, and Bruno Alicke. We acknowledge the Genentech Flow Cytometry Core group for technical assistance, including Terence Ho, George Tweet, CK Poon, Alice Tan, Cole Meyers, Kevin Le, Jonathan Paw, and Jovencio Borneo. We are grateful to the Genentech Center for Advanced Light Microscopy (CALM) core group, including Cecile Chalouni, Melissa Gonzalez Edick, Max Tauc-Adrian, David DeWitt, and Sarah Gierke, for imaging support. We thank Tamaki Jones for dosing animals in vaccination experiments and Monica Ge and Vishwa Talati for assistance with single-cell sequencing data pipelines. Finally, we acknowledge the Genentech Postdoctoral Program for its support, particularly Alexandra Colon-Rodriguez, Patsy Jaboneta, Molly Taylor, and Vishva Dixit. This study was funded by Genentech, a member of Roche.

## Author Contributions

C.B.C., C.M., and I.M. conceptualized the study. C.B.C., M.B., C.C.B., A.C.A., S.-M.L., K.L.B., G.O.-M., J.T.G., A.L., E.P., and A.R. developed or designed methodology. C.B.C, V.R. and X.P.-J. contributed to computational analysis. C.B.C., S.S., K.G.-S., L.D., C.M., and I.M. validated the findings. C.B.C., V.R., X.P.-J., C.K.C. performed formal analysis. C.B.C., M.B., C.C.B., A.C.A., S.-M.L., K.L.B., G.O.-M., J.C., I.M., C.K.C., A.R., J.T.G., A.L., E.P., and J.J. conducted the investigation. L.R., Y.L., R.B.d.S., Y.Z., S.J.T., G.A.S. S.R., S.S., K.G.-S., and L.D. provided resources, including essential study materials, reagents, and data acquisition tools. C.B.C. wrote the original draft and C.B.C., C.M., and I.M. reviewed and edited the manuscript. C.B.C. created the visualizations. C.M., I.M., S.J.T., L.R., Y.L., J.D., R.B.d.S., Y.Z., G.A.S., S.R., S.S., K.G.-S., and L.D. supervised the study. C.M. and I.M. oversaw project administration. C.M and I.M. acquired funding.

## STAR Methods

### Experimental Models and Study Participant Details

#### Mice

Mice were housed under specific-pathogen-free conditions at the Genentech animal facility. Mice were maintained in accordance with the Guide for the Care and Use of Laboratory Animals (National Research Council, 2011). Genentech is an AAALAC-accredited facility and all animal activities in this research study were conducted under protocols approved by the Genentech Institutional Animal Care and Use Committee (IACUC). Mice were housed in individually ventilated cages within animal rooms maintained under a 14 h–10 h light–dark cycle. Animal rooms were temperature and humidity controlled between 68 and 79 °F (20.0 to 26.1 °C) and from 30% to 70%, respectively, with 10 to 15 room air exchanges per hour. Male mice (aged 8– 30 weeks) that appeared healthy and free of obvious abnormalities were used for the study.C57BL/6J (JAX: 000664), C57BL/6-Tg (CAG-EGFP)1Osb/J (JAX: 003291), and B6(Cg)-Rag2tm1.1Cgn/J (JAX: 008449) mice were purchased from Jackson Laboratories. E8I.CD8A.IRES.GFP.Cre.tg_Rosa26.LSL.tdTomato.cki (GNE: 10165), XCR1.Cre.ki.B-A09-E1-1.Rosa26.LSL.tdTomato.cki_Rag2.ko (GNE: 10295), XCR1.Cre.ki.B-A09-E1-1.Rosa26.LSL.neo.YFP.cki_Rag2.ko (GNE: 11772), CD4.cre.tg.Rosa26.LSL.tdTomato.cki_OT-I.TCR.tg (GNE: 7976), Rosa26.Caags.mKate2.spliteGFP.dsDNA.tg.B6J.FO40 (GNE: 11586), CLEC9A.iCre.noneo.ki_B6.TyrC_ZBTB46.LSL.DTR.mCherry.IRES.cki (GNE: 12173), and BATF3.exon1exon2.noneo.ko.B6J (GNE: 9427) mice were bred in house.

#### Cell lines and cell culture

KPP (PDAC) cells were derived from primary tumors of male GEMM mice (KrasG12D/+;p16.p19fl/fl;Pdx1Cre) at Genentech. Parental cell lines were further engineered in house using PiggyBac vectors to overexpress different types of fluorescent reporters (mTagBFP2) or/and model antigens (decamer peptide of tumor neoantigens, including the M86 neoantigen) and/or doxycycline inducible chemokines (CXCL9, XCL1, CCL5). KPP and derivative cell lines were cultured in DMEM supplemented with 10% fetal bovine serum (FBS) and 2mM L-Glutamine at 37°C in a humidified atmosphere with 5% COLJ. Cells were maintained by passaging every 3 days, splitting at 80% confluency at a ratio of 1:10 to 1:20. For cryopreservation, cells were frozen in CryoStor10 and stored in liquid nitrogen vapor. Thawing was performed by quickly warming the vial in a 37°C water bath, followed by transfer to pre-warmed media, centrifugation to remove freezing media, and seeding into T-75 flasks. Genentech built a centralized cell bank, gCELL, to support the needs of cell-based research within Genentech. gCELL is tasked to bank verified, quality-assured cell lines for distribution throughout Genentech. This provides a consistent source of cell lines for all levels of research to enable experimental reproducibility and access to baseline information such as morphology, growth conditions. gCELL also provides an important mechanism to ensure that cell lines are used in accordance with all terms and conditions. All stocks are tested for mycoplasma before and after cells are cryopreserved. Two methods were used to avoid false-positive/negative results: Lonza Mycoalert and Stratagene Mycosensor. All of the cell lines tested negative for mycoplasma.

## Method Details

### STAMP Implantation

Experimental procedures were conducted on 8–30-week-old male mice. One day prior to microporation, ear hair was removed using Veet depilatory cream on animals under isoflurane anesthesia. For implantation, animals were anesthetized with 60–100 mg/kg ketamine and 5–10 mg/kg xylazine administered intraperitoneally. The dorsal side of the ear was immobilized with double-sided tape to expose the ventral side. Microporation was performed on the ventral side of the ear using the P.L.E.A.S.E. laser device (Pantec Biosolutions) at a depth of 71 µm, with a 9% pore density, array size of 14×14, and 2 pulses per pore at 1.4 W, 200 Hz, with a pulse length of 225µs. After microporation, 100 µl of a tumor cell suspension (KPP or derivatives) in PBS (2-4 × 10^6 cells per ear) was applied to the ventral side of the ear. The cells were incubated for 30 minutes, and then the excess suspension was removed. Microtumor growth became evident 5 to 7 days post-implantation. Tumors were checked at least twice per week once visible, with increased monitoring if adverse effects were observed, up to daily if necessary, as directed by veterinary staff. Mice were euthanized if the tumor area reached or exceeded 70% of the total ear surface, or if tumors fell outside IACUC Guidelines for Tumors in Rodents. Mice were also euthanized if ear conditions compromised normal function or if adverse effects such as severe hunching, lethargy, or moribundity were observed. Mice were weighed weekly, and those losing more than 15% body weight were weighed daily. Animals losing 20% body weight or with a body condition score of 2 or less out of 5 were euthanized or brought to the attention of the veterinary staff.

### Microscopy

To monitor tumor growth and immune infiltration, mice were imaged daily using a Leica M205 FA epifluorescence stereomicroscope equipped with a ×1.0 PlanApo lens (Leica 10450028) and an ORCAII Digital CCD camera (Hamamatsu Photonics). Epifluorescence time-lapse microscopy image series were acquired at 30X magnification over the same regions of the ear each day to monitor changes over time.

### Cell line engineering

Parental cell lines were engineered in-house using PiggyBac vectors to overexpress fluorescent reporters (mTagBFP2), model antigens (including the M86 neoantigen), and doxycycline-inducible chemokines (CXCL9, CCL5, XCL1). Cells were plated at 200,000 cells per well in a 6-well plate 12-24 hours before transfection. Transfection was performed using Lipofectamine 3000 (Thermo, Cat #: L3000015) and OptiMEM (Thermo, Cat #: 31985062), with a mixture of 1.25 µL PiggyBac Transposase Expression Vector (pBo, TaPIR id: DNA1AG47417) and 1.25 µL cargo plasmid, incubated for 15 minutes before adding to the wells. Post-transfection, cells were selected with 1-2 µg/mL puromycin (Gibco, Cat #: A1113802) or sorted for fluorescent protein expression to ensure successful integration.

### Diptheria toxin, doxycycline and Flt3L treatment *in vivo*

*In vivo* treatments were conducted on mice under protocols approved by the Genentech IACUC. Diptheria toxin (DT, Enzo Life Sciences, Cat #: BMLG1350001) was administered intraperitoneally (i.p.) at a dose of up to 500 ng per mouse to deplete cDC1 cells, with four subsequent injections at 100 ng per mouse every third day. Doxycycline (Takara Cat. #: 631311) treatment for chemokine induction involved an initial 180 µg i.p. dose on day 0, followed by 1 mg/mL Doxycycline in 5% sucrose water starting on day 0, with water bottles changed twice weekly. FLT3L-Fc was purified in house, and treatment was administered at 2 mg/kg i.p. every other day.

### T cell isolation

T cells were isolated from CD4.cre.tg Rosa26.LSL.tdTomato.cki OT-I.TCR.tg, C57BL/6-Tg (CAG-EGFP)1Osb/J (JAX: 003291), or Rosa26.Caags.mKate2.spliteGFP.dsDNA.tg.B6J.FO40 (GNE: 11586) mice. Spleens were collected and dissociated with a cell pestle into 10 ml of PBS before filtration through a 70 μm cell strainer. T cells were isolated for intravenous injection using the EasySep Mouse T Cell Isolation Kit (StemCell Technologies). A total of 4 × 10^6^ isolated T cells was injected retroorbitally per mouse.

### Tumor Dissociation

For tissue dissociation experiments, mice were euthanized and tumors were harvested. Ears were dissected into 6-well plates, and when classification by immunotype was required, tumors were individually isolated using punch biopsies and razor blades under an epifluorescence stereomicroscope. Tumors or whole ears were collected into 5 mL tubes containing 1 mL Collection Buffer (20 mL RPMI 1640, Gibco Cat No. 11875093; 100 µL FBS) and minced with scissors. Digestion Buffer (20 mL RPMI 1640, Gibco Cat No. 11875093; 80 µL collagenase P, Roche Cat. No. 43599726; 80 µL DNase, 50 mg/mL; 100 µL FBS) was added to the samples and incubated at room temperature for 15 minutes. Digestion was quenched with 0.5 mL Quenching Buffer (10 mL FBS; 20 µL eBioscience™ Protein Transport Inhibitor Cocktail, Cat No. 00-4980-03), and samples were transferred to 24-well netwell plates on ice. Tumor tissues were then mashed with cell pestles, and cell suspensions were transferred to deep well blocks for further processing. For intracellular staining, 1X eBioscience™ Protein Transport Inhibitor Cocktail was included in buffers, and cells were rested at 37°C in Culture Media (500 mL Complete RPMI with 1X Pen/Strep, 10% FBS, 1X L-Glutamax; 5 mL MEM Non-Essential Amino Acids Solution, Gibco Cat No. 11140050; 5 mL Sodium Pyruvate, Gibco Cat No. 11360070; 12.5 mL HEPES, Gibco Cat No. 15630080; 500 µL 2-Mercaptoethanol, Gibco Cat No. 31350010; 200 µL eBioscience™ Protein Transport Inhibitor Cocktail) prior to staining.

### Flow cytometry

For flow cytometry, cell suspensions were stained and analyzed as follows. Cells were resuspended PBS with Live/Dead stain (Thermo Fisher Scientific, Cat #: L23105) and incubated on ice for 30 minutes. Cells were washed twice with cold PBS and resuspended in 50 µL FACS Buffer (1X PBS, 0.5% BSA, 0.05% Na Azide) with Fc Block (1:50; BD Biosciences, Cat #553141) for 10 minutes at 4°C. Surface antigen staining was performed with the indicated antibodies (see Key Resources table). Staining was conducted for 20 minutes at room temperature in a total volume of 150 µL. For surface staining only, cells were fixed in 4% paraformaldehyde for 15 minutes on ice. After surface staining, cells were washed with cold FACS Buffer. For intracellular staining, 1X eBioscience™ Protein Transport Inhibitor Cocktail (Cat No. 00-4980-03) was included in all staining solutions prior to fixation. After surface staining, cells were fixed with 1X Foxp3 Fixation/Permeabilization working solution (eBioscience, Cat #: 00-5523-00) for 30-60 minutes at 4°C. Intracellular staining was performed overnight at 4°C in 1X Permeabilization Buffer with appropriate antibodies. Samples were analyzed using a Symphony cell analyser (BD Biosciences) or Aurora (Cytek) and analyzed using FlowJo Software (v.10.2; FlowJo).

### Adoptive transfer of antigen-matched T cells

M86 or OVA RNA-lipoplex (RNA-LPX) vaccines were assembled from M86- or SIINFEKL-coding RNAs synthesized by Genentech, using liposomes consisting of DOTMA and DOPE at a charge ratio of (+):(−) of 1.3:2.0, as described previously (Tahtinen et al Nat Immunol 2022). CD4.cre.tg Rosa26.LSL.tdTomato.cki or C57BL/6-Tg (CAG-EGFP)1Osb/J (JAX: 003291) T cell donor mice (n = 3) were vaccinated intravenously with M86 or OVA SIINFEKL RNA-lipoplex at 1, 2, and 3 weeks before T cell isolation. T cells were isolated from spleens using the EasySep Mouse T Cell Isolation Kit (StemCell Technologies). These isolated T cells were then used for adoptive transfer into Rag-2-deficient or BatF3-deficient mice. STAMP tumor implantation was performed as described, with 4 × 10^6 T cells each from M86-vaccinated, OVA-vaccinated or naive mice.

### Tissue Embedding and Cryosectioning

Tumor arrays were dissected from mouse ears, carefully positioned in Surgipath clear base molds (Ref: 3803015), and embedded in OCT compound (Tissue-Tek, Ref: 4583). Samples were snap-frozen in liquid nitrogen and then stored at -80°C until sectioning. For cryosectioning using a Leica cryostat, the OCT-embedded tissues were equilibrated to cryostat temperature (- 20 to -25°C) for 1 hour to overnight. Tissues were mounted on the cryostat chuck with additional OCT and sectioned at 10 µm thickness. A handheld 405 nm laser pen was used to visualize the mTagBFP2-expressing tumor arrays to ensure the quality of the slices and to confirm tumor presence on the tissue block and in the sections. Tissue sections were collected on Superfrost Plus Gold microscope slides (Fisherbrand, Cat No. 12-550-15) and allowed to adhere at room temperature for 5-10 minutes. Slides were then stored at -80°C or -20°C until further analysis.

### Multiplex Immunofluorescence (mIF) Staining

Mouse ear micro-tumors were processed for multiplex immunofluorescence (mIF) staining using a 3-plex Tyramide Signal Amplification (TSA) protocol on the Ventana Discovery Ultra platform (Leica). The FFPE tissue sections were first deparaffinized and rehydrated, followed by antigen retrieval using Ventana CC1 standard conditions. Primary antibodies against CD3 (SP7)(;1µg/mL, ThermoScientific,Cat#RM-9107, Clec9a (EPR2427-117) (4 µg/mL, Abcam, Cat#ab300433;;), and CD8a (21E3) (5 µg/mL, Genentech); -) were applied, with each antibody incubated for 32 minutes at 37°C. The OmniMap Rabbit HRP detection system was used for signal amplification, followed by the application of TSA reagents: Discovery Cy5 kit for CD3, Discovery Rhodamine 6G-(R6G) for Clec9a, and Akoya Opal780 kit for CD8a. Nuclear counterstaining was performed with DAPI for 16 minutes at room temperature. After staining, the sections were washed in PBS 1X and coverslipped with Prolong Gold antifade mountant. The quality of the staining was confirmed by high-resolution imaging using an Olympus VS200 slide scanner, with distinct and specific signals for CD3, Clec9a, and CD8. This mIF protocol was also validated on mouse spleen control tissue to ensure reliable and reproducible results.

### mIF Immunotype and Clustering Analysis

Tumor immunotype and immune cell spatial organization were assessed using a combination of QuPath, Python, and Fiji. To ensure accurate lesion mapping, tumors were visualized during sectioning using a handheld 405 nm laser pen, leveraging mTagBFP fluorescence to capture gross morphology. Corresponding lesions were then identified in fixed samples based on H&E and DAPI staining and defined as regions of interest (ROIs) in QuPath. T cell infiltration was evaluated manually by inspecting CD3 and CD8 fluorescence within each lesion relative to surrounding tissue, allowing for the classification of tumors as T cell-infiltrated (TC-I) or T cell-excluded (TC-Ex). Separately, cDC1 presence was assessed by Clec9a staining, determining whether lesions exhibited cDC1 infiltration (DC-In) or exclusion (DC-Ex). Based on these criteria, tumors were classified into three immunotypes: TC-In/DC-In, TC-In/DC-Ex, and TC-Ex/DC-Ex. To assess immune cell clustering, each ROI was manually inspected across DAPI, CD3, CD8, and Clec9a channels. Clustering was defined by the relative density of T cells and cDC1s within the tumor core compared to the periphery and surrounding tissue. Tumors exhibiting greater clustering of these populations within the lesion were annotated as clustered, while those with immune cells predominantly dispersed or localized to the periphery were classified as non-clustered. All analyses were performed blinded to prior classifications to ensure unbiased assessment.

### Bulk RNA Sequencing

KPP-mTagBFP2 STAMP arrays were implanted in Rag-deficient Xcr1-cre tdTomato mice with EGFP T cells. All microtumors were classified by immunotype and individually biopsied. For each replicate, five mice were grouped, with three biological replicates in total. Tumor samples were processed according to the tumor dissociation protocol described above. Cells were stained with Invitrogen LIVE/DEAD fixable dead cell stains and sorted by flow cytometry into three populations: all live/dead-negative cells, live/dead-negative tdTomato-positive cells (cDC1s), and live/dead-negative EGFP-positive cells (T cells). Sorted samples were collected into Qiagen RNeasy buffer, and RNA was isolated using the RNeasy kit following manufacturer instructions for low-input RNA. Library preparation and RNA sequencing were performed by SciGenom Labs.

### Single-Cell RNA Sequencing

KPP-mTagBFP2 STAMP arrays were implanted in Rag-deficient Xcr1-cre tdTomato mice with EGFP T cells. At endpoint, microtumors were individually biopsied, classified by immunotype, and processed according to the tumor dissociation protocol described above. To exclude non-viable cells, samples were first stained with LIVE/DEAD Fixable Dead Cell Stain (Invitrogen, Cat #L23105). To minimize nonspecific antibody binding, cells were resuspended in 50 µL of FACS buffer (PBS + 2% BSA + 0.01% Tween-20) per sample and incubated with TruStain FcX (anti-mouse CD16/32, BioLegend, Cat #101320). For sample multiplexing, TotalSeq-A hashing antibodies (2 µL per 1 × 10L cells per sample, BioLegend) were applied in a total staining volume of 100 µL, with each sample receiving a unique hashing antibody for downstream demultiplexing. For surface marker identification, cells were stained with fluorescently conjugated antibodies targeting CD45 (BUV395), MHCII (AF700), CD11c (APC-Cy7) for dendritic cells, and CD90 (FITC) for T cells to enrich immune cell subsets of interest. Where applicable, CITE-seq antibodies (4.5 µL per 1 × 10L cells per sample, BioLegend TotalSeq-A) were included for protein-level characterization via sequencing-based barcoding. Following staining, cells were washed three times with PBS + 2% BSA + 0.01% Tween-20, resuspended in PBS + 2% BSA, counted using a hemocytometer with Trypan blue exclusion, and passed through a 40 µm strainer to remove cell aggregates.

Samples were sorted on a BD FACS Aria, and sorted populations were manually counted. To ensure unbiased representation, samples were pooled in equal proportions across immunotypes. A total of 96,000 CD45L tumor-infiltrating cells, 20,000 CD45LMHCIILCD11cL tumor-infiltrating cDC1s, and 20,000 CD45LCD90L T cells were loaded onto a 10X Chromium Controller for single-cell partitioning into gel bead-in-emulsion (GEM) droplets. cDNA libraries were prepared using the 10X Genomics 5’ Gene Expression Kit, with additional libraries constructed for hashtag oligonucleotides (HTOs) and antibody-derived tags (ADT). During 10X library preparation, cDNA amplification was modified to incorporate HTO and ADT primers. Libraries were sequenced at a targeted depth of 20,000 reads per cell.

## Quantification and Statistical Analysis

## Key Resources Table

**Supplementary Figure 1.**
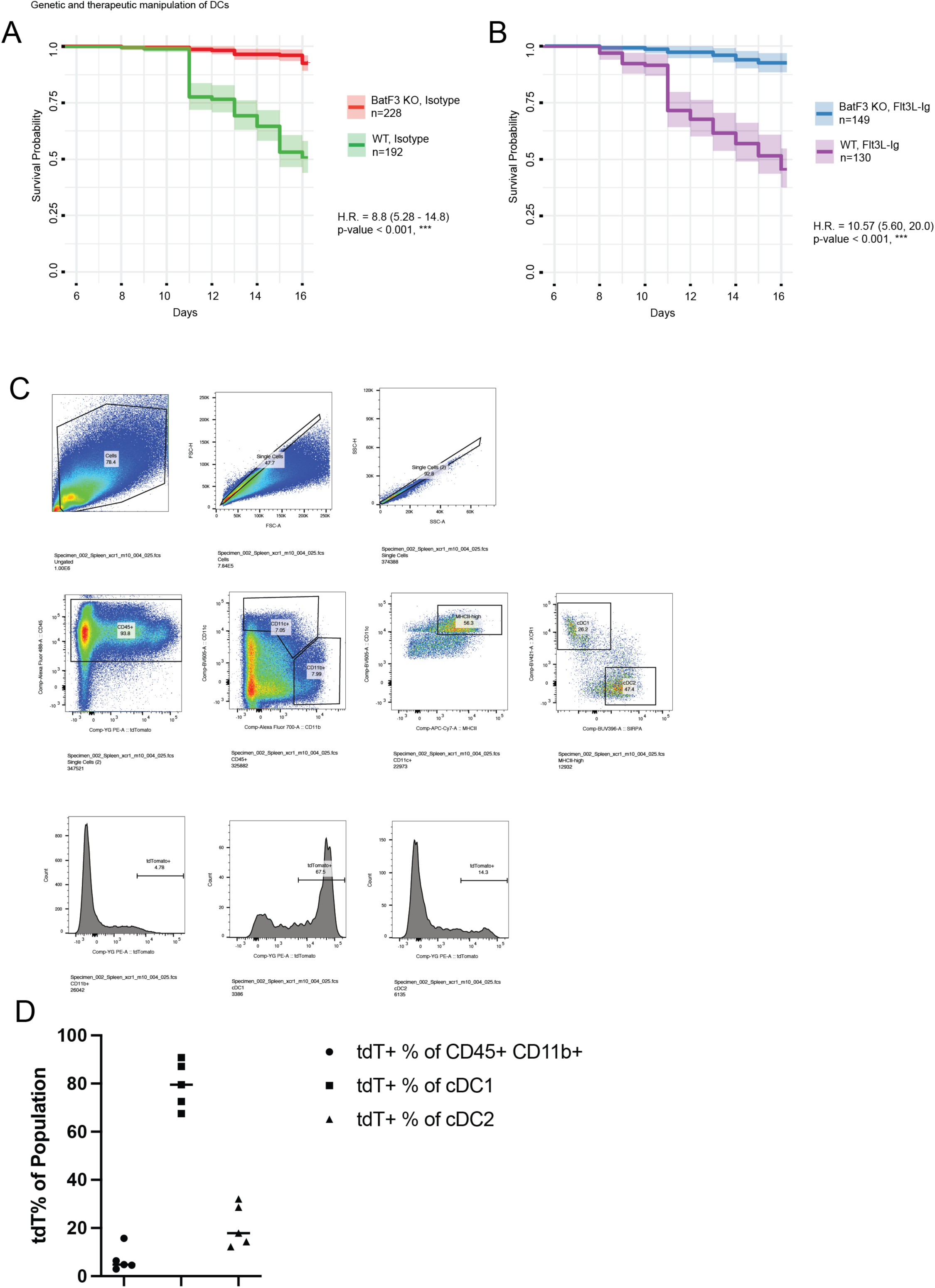
cDC1 and T cell localization defines distinct immunotypes in mouse and human tumors. (A) Kaplan-Meier curves showing individual microtumor rejection in KPP-mTagBFP2 STAMP arrays implanted in BatF3-deficient or WT animals. *n* = 5 animals per group. (B) Kaplan-Meier curves showing individual microtumor rejection in KPP-mTagBFP2 STAMP arrays implanted in BatF3-deficient or WT animals treated with Flt3L. *n* = 5 animals per group. (C) Gating strategy for identifying tdTomato+ cells in Xcr1-Cre; Rosa26-LSL-tdTomato; Rag2-deficient animals reconstituted with EGFP+ T cells. Data shown for one representative animal. (D) Percentage of tdTomato+ cells in CD45+CD11b+, cDC1, or cDC2 populations as defined by the gating strategy in (C). *n* = 5 animals.

**Supplementary Video 1. Calcium-Flashing Activity in a Representative TC-In/DC-In Tumor**

Time-lapse video of calcium-flashing activity in a representative T cell-inflamed, DC-Innflamed (TC-In/DC-In) tumor from a KPP-mTagBFP2 STAMP array. Tumor cells express the GCaMP6 calcium sensor, which fluoresces upon calcium influx as a proxy for T cell-induced pore formation, indicating cytolytic activity. The tumor was implanted into a Rag2-deficient Xcr1-Cre Rosa-tdTomato mouse reconstituted with mKate+ T cells. GCaMP6 fluorescence (green) shows active tumor cell killing, and flashing intensity correlates with the level of T cell cytotoxicity observed. Scale bar = 250 µm.

**Supplementary Video 2. Calcium-Flashing Activity in a Representative TC-In/DC-Ex Tumor**

Time-lapse video of calcium-flashing activity in a representative T cell-inflamed, DC-excluded (TC-In/DC-Ex) tumor from a KPP-mTagBFP2 STAMP array. Tumor cells express the GCaMP6 calcium sensor, which fluoresces upon calcium influx as a proxy for T cell-induced pore formation, indicating cytolytic activity. The tumor was implanted into a Rag2-deficient Xcr1-Cre Rosa-tdTomato mouse reconstituted with mKate+ T cells. GCaMP6 fluorescence (green) shows lower flashing intensity compared to the TC-In/DC-In tumor, reflecting reduced T cell cytotoxic activity in the absence of cDC1s. Scale bar = 250 µm.

**Supplementary Figure 2.**
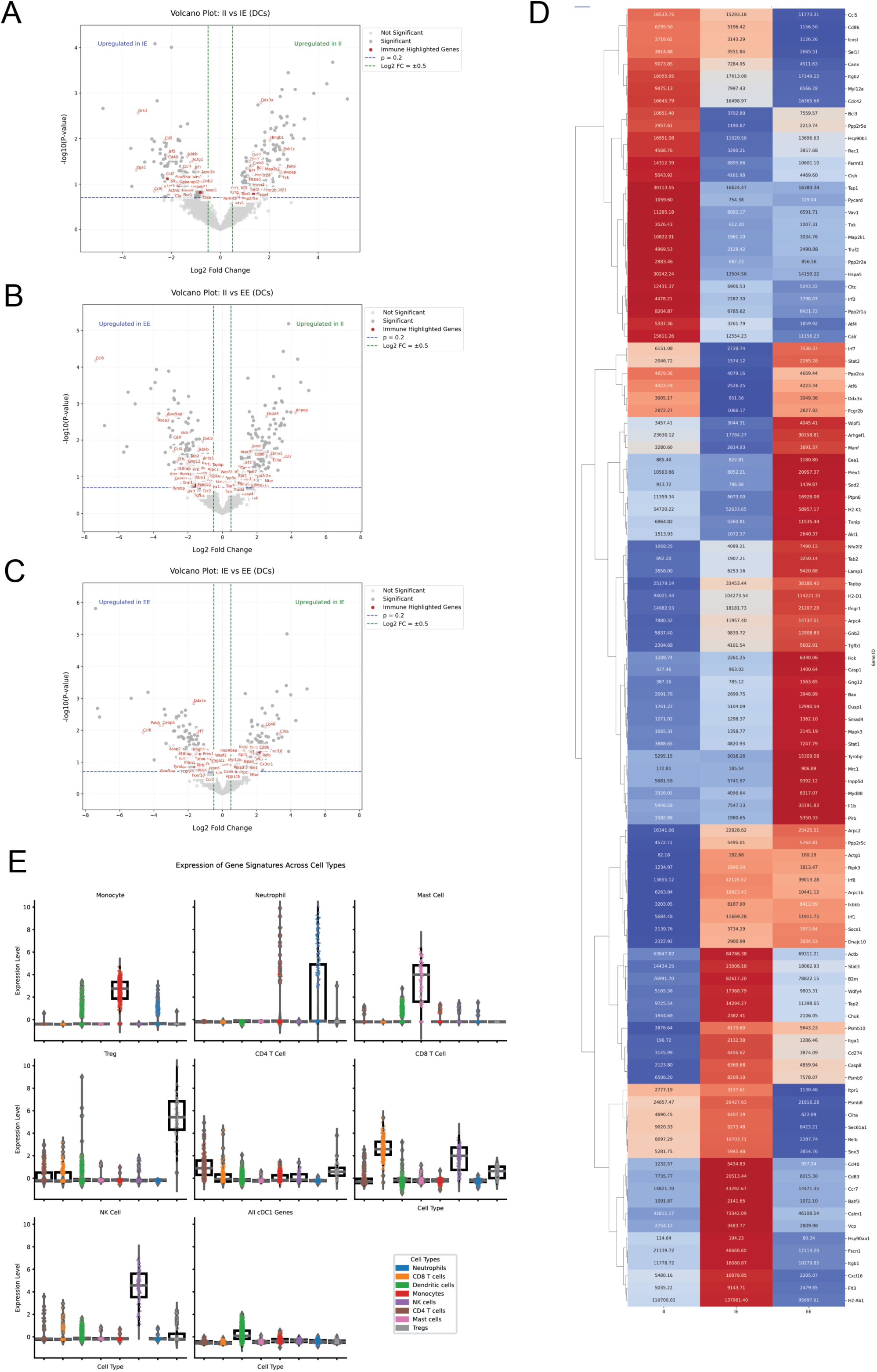
Transcriptional Profiles of Dendritic Cells Across Tumor Immunotypes. (A–C) Volcano plots showing differentially expressed genes (DEGs) in dendritic cells (cDC1s) isolated from tumors classified by immunotype. (A) TC-In/DC-In (T cell-inflamed, DC-Innflamed) versus TC-In/DC-Ex (T cell-inflamed, DC-excluded). (B) TC-In/DC-In versus TC-Ex/DC-Ex (T cell-excluded, DC-excluded). (C) TC-In/DC-Ex versus TC-Ex/DC-Ex. DEGs were identified using bulk RNA sequencing of Xcr1+ cDC1s from KPP-mTagBFP2 STAMP tumor arrays implanted in Xcr1-Cre; Rosa26-LSL-tdTomato Rag-deficient mice reconstituted with GFP+ T cells. Data represent pooled tumors from 5 mice per immunotype, with three replicates. (D) Heatmap of differentially expressed immune-related genes across all three immunotypes (TC-In/DC-In, TC-In/DC-Ex, and TC-Ex/DC-Ex), highlighting genes associated with activation, migration, antigen presentation, and stress response. Data are normalized and hierarchically clustered.

**Supplementary Figure 3.**
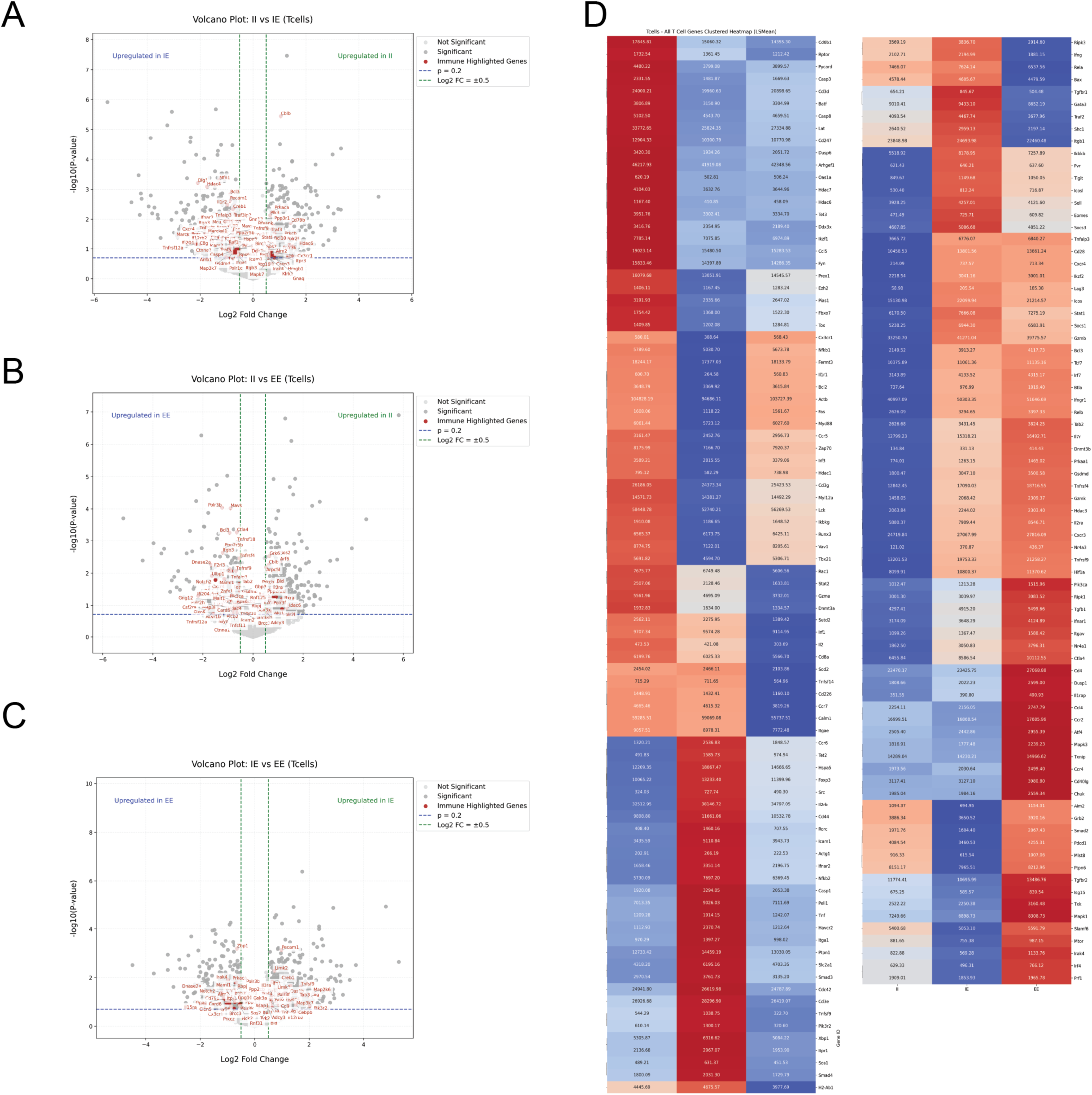
Transcriptional Profiles of T Cells Across Tumor Immunotypes. (A–C) Volcano plots showing differentially expressed genes (DEGs) in T cells isolated from tumors classified by immunotype. (A) TC-In/DC-In (T cell-inflamed, DC-Innflamed) versus TC-In/DC-Ex (T cell-inflamed, DC-excluded). (B) TC-In/DC-In versus TC-Ex/DC-Ex (T cell-excluded, DC-excluded). (C) TC-In/DC-Ex versus TC-Ex/DC-Ex. DEGs were identified using bulk RNA sequencing of T cells from KPP-mTagBFP2 STAMP tumor arrays implanted in Xcr1-Cre; Rosa26-LSL-tdTomato Rag-deficient mice reconstituted with GFP+ T cells. Data represent pooled tumors from 5 mice per immunotype, with three replicates. (D) Heatmap of differentially expressed immune-related genes across all three immunotypes (TC-In/DC-In, TC-In/DC-Ex, and TC-Ex/DC-Ex), highlighting genes associated with cytotoxicity, activation, exhaustion, and regulatory functions. Data are normalized and clustered by gene expression profiles.

**Supplementary Figure 4.**
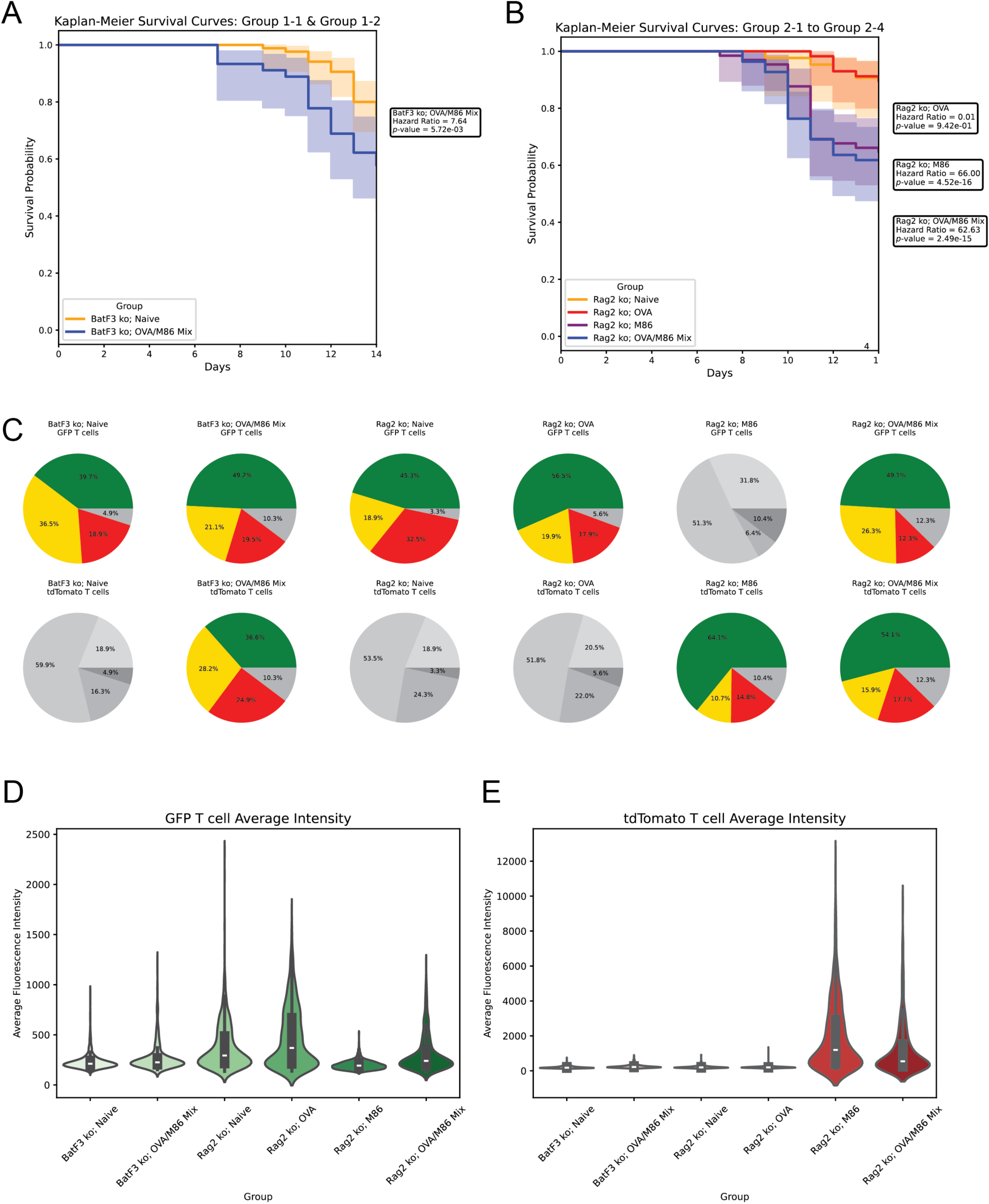
cDC1s Enhance Tumor Rejection and T Cell Recruitment in the STAMP Model. (A) Kaplan-Meier survival analysis of BatF3 KO mice implanted with KPP-mTagBFP2-M86 STAMP microtumor arrays and receiving either naïve GFP+ T cells or a mixed transfer of OVA vaccine elicited GFP+ T cells and M86 vaccine elicited tdTomato+ T cells (n=5 mice per group). (B) Kaplan-Meier survival analysis of Rag2 KO mice implanted with KPP-mTagBFP2-M86 STAMP microtumor arrays and receiving either naïve GFP+ T cells, OVA vaccine elicited GFP+ T cells, M86 vaccine elicited tdTomato+ T cells, or a mix of OVA and M86 vaccine elicited T cells (n=5 mice per group). (C) Immunotype distribution for GFP+ and tdTomato+ T cells across tumor regions. Pie charts indicate the proportion of inflamed (green), excluded (red), and desert (yellow) tumors for each T cell population. Grayed-out pie charts indicate conditions where the respective T cell population was not present (e.g., tdTomato+ proportions in conditions without tdTomato+ T cells). (D–E) Quantification of average fluorescence intensity for GFP+ T cells (D) and tdTomato+ T cells (E) across tumors, reflecting T cell infiltration and retention. Data from the same experiment as (A–C), n=5 mice per group.

